# Synergistic Material-Topography Combinations to Achieve Immunomodulatory Osteogenic Biomaterials

**DOI:** 10.1101/2020.04.29.067421

**Authors:** Laurence Burroughs, Mahetab H. Amer, Matthew Vassey, Britta Koch, Grazziela P Figueredo, Blessing Mukonoweshuro, Paulius Mikulskis, Aliaksei Vasilevich, Steven Vermeulen, Ian L. Dryden, David A. Winkler, Amir M. Ghaemmaghami, Felicity R. A. J. Rose, Jan de Boer, Morgan R. Alexander

## Abstract

Human mesenchymal stem cells (hMSCs) are widely represented in ongoing regenerative medicine clinical trials due to their ease of autologous implantation. In bone regeneration, crosstalk between macrophages and hMSCs is critical with macrophages playing a key role in the recruitment and differentiation of hMSCs. However, engineered biomaterials able to both direct hMSC fate and modulate macrophage phenotype have not yet been identified. A novel combinatorial chemistry-microtopography screening platform, the *ChemoTopoChip*, is used to identify materials suitable for bone regeneration by screening with human immortalized mesenchymal stem cells (hiMSCs) and human macrophages. The osteoinduction achieved in hiMSCs cultured on the “hit” materials in basal media is comparable to that seen when cells are cultured in osteogenic media, illustrating that these materials offer a materials-induced alternative in bone-regenerative applications. These also exhibit immunomodulatory effects, concurrently polarizing macrophages towards a pro-healing phenotype. Control of cell response is achieved when both chemistry and topography are recruited to instruct the required cell phenotype, combining synergistically. The large library of materials reveals that the relative roles of microtopography and material chemistry are similar, and machine learning identifies key material and topographical features for cell-instruction.

## Main Section

Bone repair is a complex yet highly organized process involving interactions between multiple cell types, molecular signals, and interactions with the extracellular environment.^[1]^ Currently, autologous bone grafts remain the gold standard in bone regeneration because of their osteogenicity, osteoinductivity, osteoconduction and osteointegration characteristics.^[2],[3]^ Synthetic bone substitutes, such as calcium phosphate (CaP) ceramics, have proven safe and biocompatible but often lack the osteogenicity needed to support bone healing.^[4]^ There is increased interest in the use of hMSCs in combination with synthetic biomaterials to provide a potential way of overcoming these challenges in autologous bone grafting.^[5], [6]^

The inherent multipotency of hMSC cells has allowed *in vitro* cultured models to be used, in combination with synthetic biomaterials, to differentiate cells into osteoblasts without osteogenic supplements. They have also been shown to form bone *in vivo*, a process driven by topography, protein adsorption to surface chemistry and phosphorus delivery to the cells.^[5], [6], [7], [8], [9]^ Furthermore, there is growing evidence demonstrating a reciprocal functional role of macrophage polarization in hMSC osteoblast differentiation, with pro-healing M2 macrophages enhancing hMSC osteoblastic differentiation.^[10]^ Hence, the crosstalk between macrophages and MSCs plays a key role in normal bone repair.^[11], [12], [13]^

The ability to tune or modulate the foreign-body response to a biomaterial is an ongoing challenge in the field of regenerative medicine.^[14]^ This is further complicated when designing materials for tasks such as induction of osteogenic differentiation of hMSCs at the site of implantation. Surface chemistry^[15], [16]^ and surface topography^[17], [18], [19]^ have both been shown to enhance differentiation and proliferation of stem cells yet, to our knowledge, no biomaterials have been reported capable of simultaneously directing differentiation of hMSCs and polarizing macrophages towards an M2 state.

To enable a wide range of chemistries and topographies to be screened, a combinatorial high-throughput screening tool, the *ChemoTopoChip*, was developed. This platform contains 980 microtopography and materials chemistry combinations that can simultaneously probe the combined effects on cellular response. The screening platform was used to identify materials that can direct hiMSC differentiation towards an osteoblastic lineage and that polarize human macrophages towards a pro-healing M2 phenotype, two cell phenotypes that have regenerative medicine applications. The design comprised 36 Topo units of size 500 × 500 µm, including one flat control (**Figure 1 b**-**c**), arranged in 3 × 3 mm ChemoTopo units. These are repeated 28 times, each with a different chemical functionalization (**Figure 1 a**). The microtopographies used were chosen from previous TopoChip screens to maximize the morphological differences of MSCs (see **Figure S1** for high-resolution images).^[20]^ The chemistries were chosen from libraries of (meth)acrylate and (meth)acylamide monomers to provide maximum chemical diversity (see **Figure S2** for structures). The monomers are used to functionalize the surface of topographically molded chips, which minimizes differences in material compliance between chemistries sensed by the attached cells (see **Table S1** for AFM modulus measurements).

**Figure 1.**
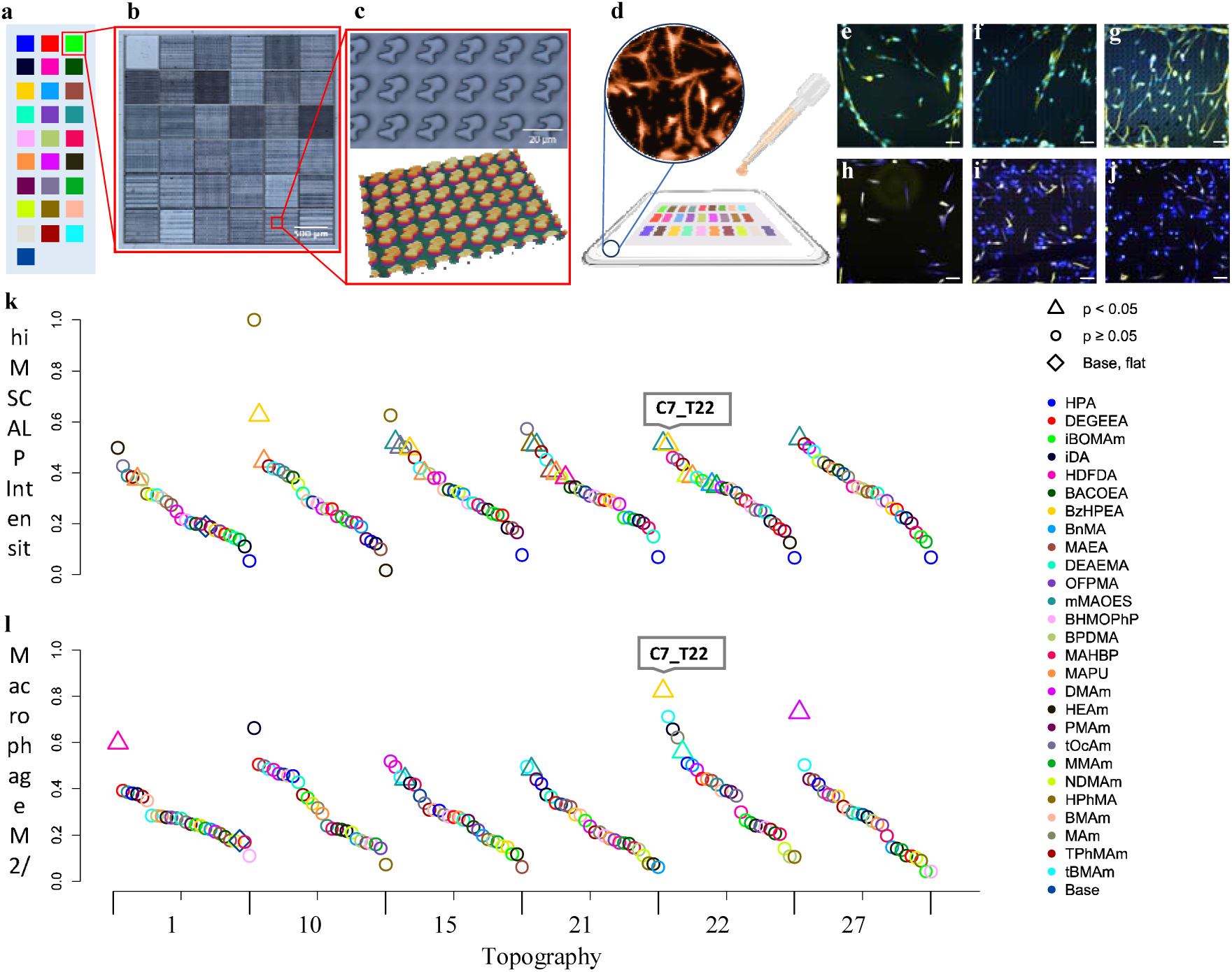
**a)** Schematic showing ChemoTopoChip layout; **b)** Interference profilometer imaged ChemoTopo unit (30 µm high walls separate each Topo unit); **c)** Example features from a ChemoTopo unit; **d)** Cells seeded onto ChemoTopoChips **e)** hiMSCs on flat TMPMP-co-TEGDA area (blue = ALP, yellow = α-tubulin); **f)** hiMSCs on TMPMP-co-TEGDA + Topo 3 area (blue = ALP, yellow = α-tubulin); **g)** hiMSCs on mMAOES + Topo 3 area (blue = ALP, yellow = α-tubulin); **h)** Macrophages on flat TMPMP-co-TEGDA area (blue = IL-10, yellow = TNFα); **i)** Macrophages on TMPMP-co-TEGDA + Topo 22 area (blue = IL-10, yellow = TNFα); **j)** Macrophages on BzHPEA + Topo 22 area (blue = IL-10, yellow = TNFα); **k)** Rank ordered ordered hiMSC cell number across the ChemoTopoChip (N = 3, n = 3) on selected topographies; **l)** Rank hiMSC ALP intensities across the ChemoTopoChip (N = 3, n = 3) on selected topographies; **m)** Rank ordered human macrophage cell number across the ChemoTopoChip (N = 2, n = 3) on selected topographies; **n)** Rank ordered human macrophage M2/M1 ratio across the ChemoTopoChip (N = 2, n = 3) on selected topographies

A silicon mold was fabricated from the ChemoTopoChip design using photolithography and etching to produce the negative master of the topographies. The desired features were produced from this master by injecting a 1:2 mixture of trimethylolpropane tri(3-mercaptopropionate):tetra(ethylene glycol) diacrylate (1:2 TMPMP:TEGDA) monomers containing the photoinitiator 2,2-dimethoxy-2-phenylacetophenone (DMPA) between a methacrylate-functionalized glass slide and the silicon master. UV curing and solvent washing then provided the molded ChemoTopoChip substrate, chosen because similar photopolymerized thiol-ene systems have been reported as tough shape memory, flexible materials offering low shrinkage stress that are sufficiently transparent to allow transmission optical imaging.^[21]^ Functionalization of the ChemoTopo units was carried out by deposition of 50% w/v or 75% v/v monomer solutions in *N,N*-dimethylformamide (DMF) containing 0.05% w/v DMPA onto each ChemoTopo unit; further UV curing and washing steps delivered the final ChemoTopoChip (**Figure 1 a**-**c, Figure S3, Figure S4**).

To first investigate the osteoinductive potential of the materials, hiMSCs were seeded on 3 replicate chips in 3 independent experiments (N = 3, n = 3) and cultured in basal culture media (a non-selective, undefined culture medium containing all elements required for cell growth but not containing osteo-inductive supplements to induce stem cell differentiation). After 5 days, samples were fixed and stained for both α-tubulin (cytoskeletal marker) and alkaline phosphatase (ALP, an early osteogenic marker), and analyzed using an automated high-throughput fluorescence microscope. ALP expression is a widely used osteogenesis marker as it is known to be involved in bone formation, plays an essential role in matrix mineralization and is induced by a range of osteogenic molecules.^[22]^ Images were processed using CellProfiler software^[23]^ to quantify cell number and ALP staining intensity on each individual topography-material combination. The ALP staining intensity and cell number were normalized to that of the flat TMPMP-co-TEGDA Topo unit within each ChemoTopoChip sample.

A diverse range of hiMSC cell morphologies and attached cell numbers could be seen across the ChemoTopoChip (**Figure 1 e**-**g, Figure S4** and **S5**), with cells displaying an elongated shape and alignment to the topographies on some ChemoTopo units (e.g. **Figure 1 f, Figure S5** and **S6** E2, E3) in contrast to more uniform cell spreading and random alignment on others (eg. **Figure 1 e, Figure S5** and **S6** B3, C4). Previous TopoChip screens using hMSCs revealed a similar range of cell morphological responses, where more elongated cells were linked to ALP upregulation.^[14]^ To explore the range and identify trends it is useful to plot all the results as both heatmaps and to rank order all results to see the range for all ChemoTopo units (see **Figure S7** and **Figure S8 a-b**). For an exploratory method we indicate the combinations which were determined to have p-value < 0.05 from a two independent sample equal variance t-test, although we note that the sample sizes are small here and this does not account for the *multiple comparisons problem*. Analysis of the mean integrated ALP expression per cell for each topography-material combination showed that 113 exhibited the most significant upregulation of this osteogenic marker (p < 0.05) compared to the flat base Topo unit (see **Table S2** for full list), with all of these displaying a higher ALP intensity than the flat base material region used as a control comparator. Visual inspection reveals trends across various chemistries, e.g. monomers 12 (mono-2-(methacryloyloxy)ethyl succinate, mMAOES) and 20 (*N*-tert-octylacrylamide, tOcAm), suggesting that a group of chemistries induce upregulation of ALP intensity relative to the mean; equivalent topographical trends were less evident indicating that micro-topographical stimuli did not dominate across the range of chemistries used. A total of 103 combinations were found to have higher normalized cell number than the flat base region (p < 0.05, total area taken into account), but none lower (**Table S2** for full list). All combinations containing microtopographies showed greater cell numbers than those of chemistries on flat surfaces (Topo 1), suggesting that topography was also a driver for hiMSC attachment.

The mean ALP fluorescence intensity per cell of the hiMSC positive control was compared to that of the ChemoTopoChip topography-material combinations. The positive control consists of hiMSCs stimulated to differentiate down the osteogenic lineage by culturing in osteo-inductive media, in contrast to the basal media used in the ChemoTopoChip experiments (supplements detailed in **Supporting Information**). No difference in ALP upregulation was observed between the top 50 ChemoTopoChip ALP hits and the positive control cultured in osteogenic media (p < 0.001, see **Figure S9**). These materials therefore induce a similar osteogenic state of the cells, as measured by ALP upregulation, to that of osteo-inductive supplements typically used to differentiate hMSCs to osteoblasts.

Immunomodulation was next screened by seeding primary human monocytes onto ChemoTopoChips for 6 days followed by testing for differentiation into macrophages and polarization to the M1 or M2 phenotype. Monocytes were isolated from peripheral blood of two independent donors, with 3 replicates carried out for each. To determine the polarization status of the cells, samples were fixed and stained for intracellular expression of the pro- and anti-inflammatory cytokines tumor necrosis factor α (TNFα, M1 polarization indicator) and interleukin-10 (IL-10, M2 polarization indicator) respectively, and analyzed using high-throughput fluorescence microscopy. Images were processed using CellProfiler software^[19]^ with an image analysis pipeline designed to quantify cell attachment using DAPI nuclear staining and mean fluorescence intensity (MFI) across each Topo unit for the IL-10 and TNFα channels. The IL-10 and TNFα MFI and cell number were normalized to the values from the flat base TMPMP-co-TEGDA Topo unit. The ratio of M2/M1 cells was taken to be the ratio of the IL-10/TNFα MFIs.

Plotting the normalized macrophage cell number and M2/M1 ratio as a ranked scatter plot ordered by topography (**Figure S8 c-d**) and as heatmaps (**Figure S10**) indicated that chemistry has a greater influence over human macrophage polarization than topography. This is due to the dominance of chemistries 2-(4-benzoyl-3-hydroxyphenoxy)ethyl acrylate (BzHPEA), *N,N*’-Dimethylacrylamide (DMAm) and heptadecafluorodecyl acrylate (HDFDA) in the highest scoring combinations. Similar ranges of responses were observed for all topography-material combinations, including those containing flat areas. As was seen with hiMSC attachment, the range of normalized cell number for combinations containing microtopographies was greater than those for flat areas; topography is therefore also important for macrophage attachment. Visual inspection identified some topography-material combinations that were hits for both data sets, with BzHPEA (chemistry 7) in combination with topography 22 appearing strongest for both ALP upregulation in hiMSCs and M2 polarization in macrophages (**Figure 1 k-l**).

For analysis of the ChemoTopoChip data, unfunctionalized TMPMP-co-TEGDA molded topographies and flat area chemistries were used as control chemistries and topographies. Hit topography-material combinations were compared to these controls to assess their synergy ratios (*SR*) (see **Supporting Information** for method). For the hiMSC data set, 15 of the 103 hit combinations that showed greater cell numbers than the flat TPMP-co-TEGDA control were determined to be synergistic (**Figure 2 a**, see **Table S2** for full analysis of all synergistic combinations); additionally, 2 of the 113 hit combinations directing osteogenic differentiation were determined to be synergistic (**Figure 2 b**, see **Table S2** for full analysis of all synergistic combinations). For the hiMSC cell number, 2 combinations appeared to be antagonistic (**Figure 2 a**). For the human macrophage cell number, 2 topography-material combinations exhibited a synergistic effect (**Figure 2 c**, see **Table S3** for full analysis). Of the top 22 human macrophage polarization combinations, 4 were determined to be synergistic (**Figure 2 d**, see **Table S3** for full analysis). A total of 3 combinations were found to promote upregulation of ALP in hiMSCs *and* polarize human macrophages towards an M2 phenotype (**Figure 2 b, d**-**f**). The top hit, BzHPEA in combination with Topo 22, was found to be synergistic for both datasets (**Figure 2 b, d**).

**Figure 2.**
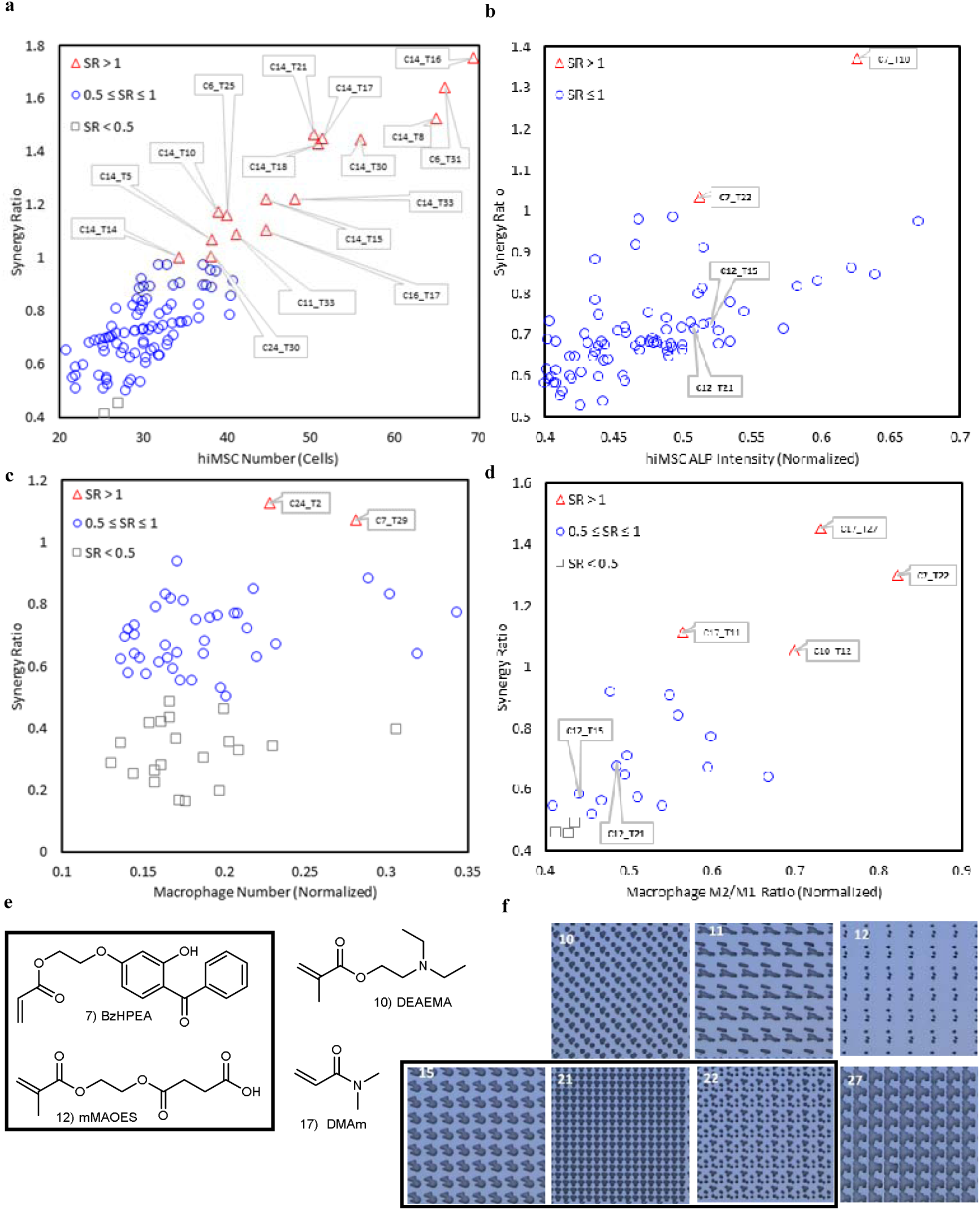
**a)** *SR* plotted versus hiMSC cell number; **b)** *SR* plotted versus hiMSC ALP intensity (normalized); **c)** *SR* plotted versus human macrophage normalized cell number; **d)** *SR* plotted versus human macrophage M2/M1 ratio (normalized); **e)** Selected hit chemistries from macrophage and hiMSC datasets (see **Figure S2** for full list of chemistries). Coincident M2/M1 and ALP hits highlighted; **f)** Selected hit topographies from macrophage and hiMSC datasets (see **Figure S1** for full list of topographies). Coincident M2/M1 and ALP hits highlighted

To investigate the feasibility of extracting rules from ChemoTopoChip screening data to inform future materials development, we used machine learning (ML) methods to generate quantitaive structure-activity relationships. A combination of chemistry descriptors, “1-hot” binary variables indicating the presence or absence of a chemistry in any given combination, and topographical shape descriptors generated from CellProfiler^[19]^ was used to model both data sets using the Random Forest algorithm^[24]^ (**Figure 3**). The 1-hot encoding method generates a vector with length equal to the number of categories in the data set. If a particular category is present, that category’s position is set to 1 and all other positions in the vector are zero. Thus, 1-hot encoding is a process for converting categorical variables into a form that machine learning algorithms can use to generate predictive models.

**Figure 3.**
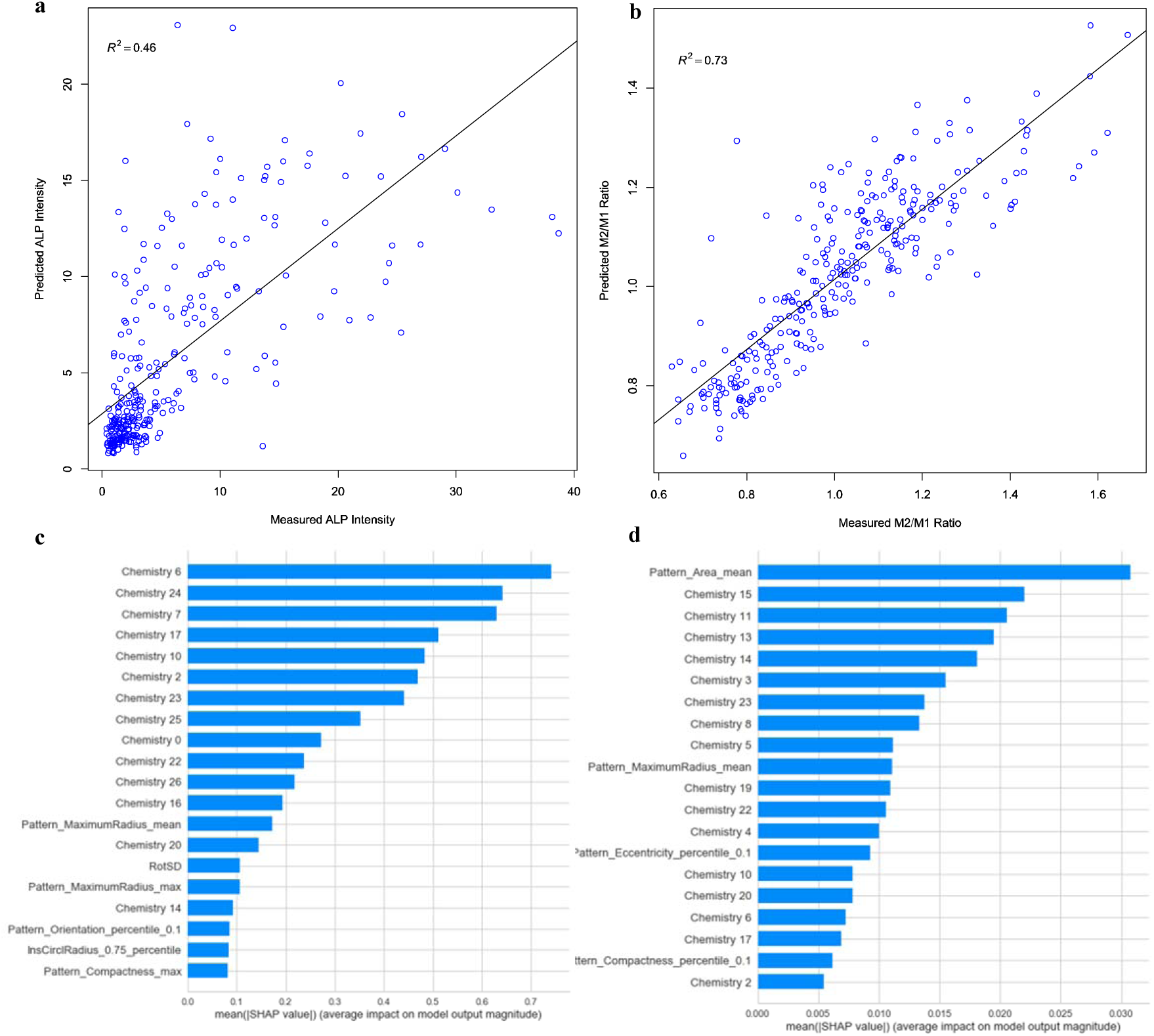
**a)** hiMSC ALP intensity random forest model using indicator variables for chemistries and topographical descriptors; **b)** human macrophage polarization random forest model using indicator variables for chemistries and topographical descriptors; **c)** hiMSC ALP intensity random forest model top contributions; **d)** human macrophage polarization random forest model top contributions

The macrophage M2/M1 ratio model had a strong correlation between the ML-predicted and observed values (*R*^2^ = 0.73). The size of the topographical features was identified as being important for macrophage polarization. Features with mean areas < 50 µm^2^ and maximum radii of 1-3 µm generated highest M2/M1 ratio (see **Figure S11** for polarization vs. descriptor correlations). The circularity of the topographical features was a strong contributor to the model, with smaller eccentricities producing the greatest increase in macrophage M2 polarization. Topographical descriptors had a greater impact on the M2/M1 human macrophage model than on the hiMSC ALP intensity model (i.e. topography plays a larger role in macrophage polarization than in hiMSC osteoinduction). This is consistent with the phagocytic nature of macrophage cells, which are designed to engulf bacterial cells and small particles. The correlation with shape may also imply that features of particles that are closer to being circular (which is highly unlikely in nature) trigger more M2 cytokines, whereas higher eccentricity (as observed with most microorganisms and foreign particles) trigger M1 response in line with the body’s natural defenses. These analyses illustrate the potential of the ChemoTopoChip and ML for uncovering complex relationships between topography, chemistry, and cell response that offer opportunities for bespoke cell phenotype control using materials design alone.

The hiMSC ALP intensity Random Forest model produced a relatively low correlation between predicted and observed ALP induction (*R*^2^ = 0.46). Difficulties in modelling stem cell responses in polymeric biomaterials has been noted previously,^[13]^ in that case due to a relatively small number of polymers with diverse chemotypes driving desirable cell responses. There were therefore insufficient examples of each chemical feature for the ML models to generate rules from. Topographical descriptors identified as being important in the hiMSC ALP model included the size of the topographical features (see **Table S4** for list of feature descriptions), with features ≤ 3.5 µm radius increasing ALP expression. However, this trend was not as strong as that observed for macrophage polarization (see **Figure S12** for ALP upregulation vs. descriptor correlations). Orientation of topographical features also contributed to the model, with those having a small number (< 10%) of features rotated > 25° relative to the x-axis of the Topo unit walls driving an increase in ALP expression.

In previous modelling studies of biological responses to polymer libraries, signature and other fragment-based molecular descriptors and traditional Dragon molecular descriptors have been shown to represent surface chemistries well. These descriptors generated robust, predictive models for diverse biological responses.^[25]^ Paradoxically, in the current study, these types of chemical descriptors were unable to generate ML models for the ChemoTopoChip data that were as accurate as the models using simple 1-hot descriptors to encode the identities of the polymer chemistries. We propose that this is due to the 28 chemistries on the ChemoTopoChip being designed to have very diverse chemistries, in order to cover chemical space as widely as possible (albeit sparsely). The key chemical fragments and resultant descriptors are also therefore very sparse. ML models cannot learn features that are not sufficiently represented in the data set, hence the combination of high chemical diversity and low number of samples resulted in inadequate information on which to train the ML models, resulting in lower prediction accuracies.

In summary, we have employed the novel ChemoTopoChip screening platform for biomaterials discovery, harnessing the potential of both chemistry and topography, to develop immunomodulatory materials suitable for use in bone regenerative applications. Analysis of the hiMSC and human macrophage datasets has identified a range of novel chemistry-topography combinations that surpass the material-instructive cues provided by either alone, with BzHPEA in combination with Topo 22 being synergistic for both cell types. Numbers of each cell type and hiMSC ALP upregulation spanned similar ranges for the chemistries and microtopographies studied, but macrophage polarization was more strongly influenced by chemistry. Thus, chemical and topographical features are both important drivers when designing biomaterials for simultaneous control of multiple cell types. Modelling of the human macrophage polarization data showed that small, cylindrical pillars of < 3 µm radius directed macrophage polarization towards an anti-inflammatory phenotype. The size and orientation of topographical features was also important for hiMSC ALP expression, with features of ≤ 3.5 µm radius and rotation of > 25° relative to the x-axis of the Topo unit walls providing strongest upregulation of ALP. Data generated by the ChemoTopoChip has been shown to be very amenable to machine learning methods, facilitating the development of models that aid design and discovery of bio-instructive materials.

## Acknowledgements

This research is supported by funding from the Engineering Physical Sciences Research Council (EPSRC) under the Programme Grant Next Generation Biomaterials Discovery EP/N006615/1. AV, SV and JdB are funded by the European Union’s Seventh Framework Programme (FP7/2007-2013) under grant agreement no 289720. AV acknowledge the financial contribution of the Province of Limburg. SV acknowledges the financial support of the European Union’s Horizon 2020 Programme (H2020-MSCA-ITN-2015; Grant agreement 676338). The authors would like to thank Dr Chris Gell (School of Life Sciences Imaging and Microscopy (SLIM) Facility) for assistance with high throughput microscopy.

## Disclosure of potential financial conflicts

Jan deBoer is a founder and shareholder of Materiomics b.v.

---

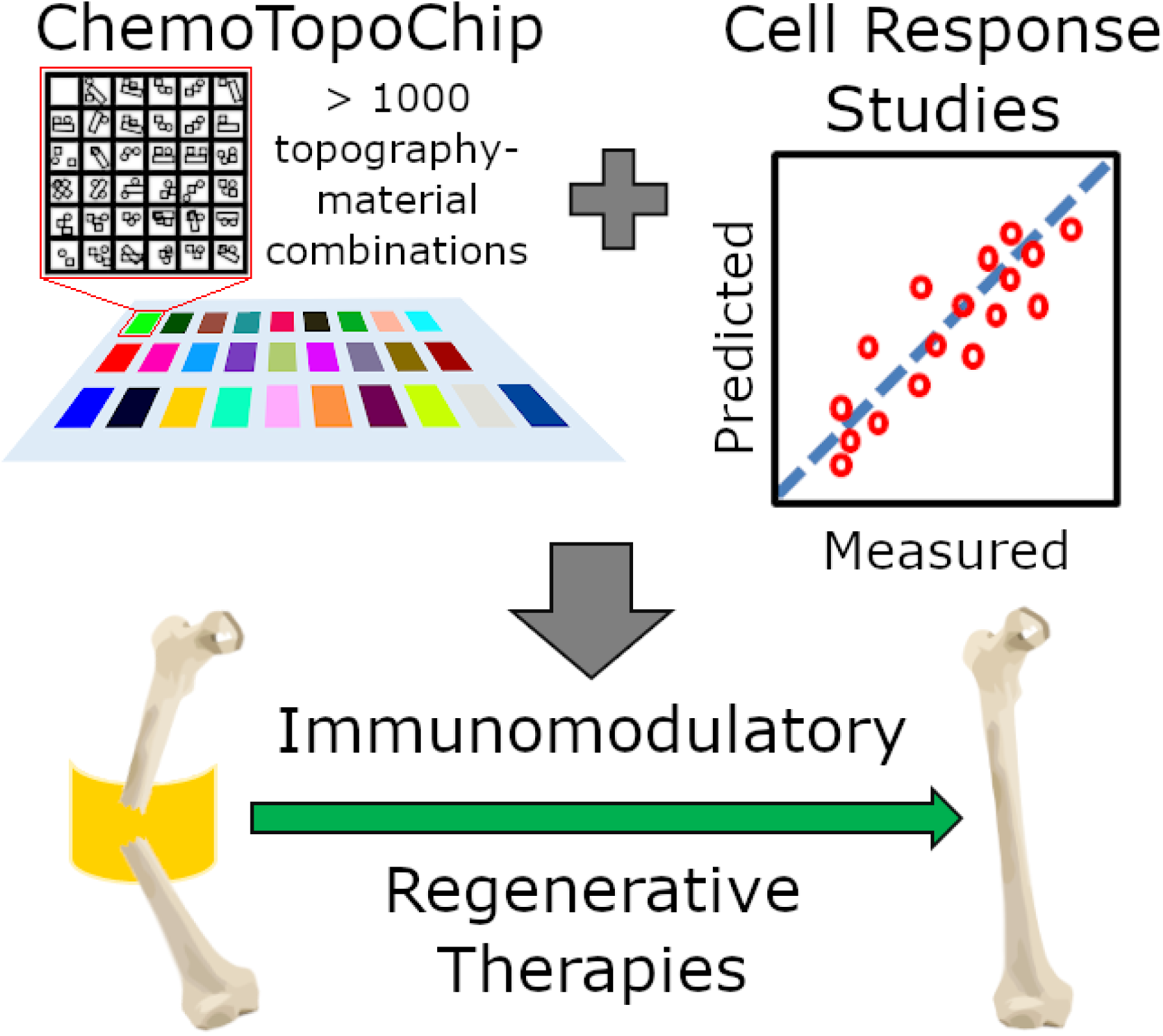

## Supporting Information

### Methacrylate Functionalization of Glass Slides

Glass slides (26 mm × 50 mm × 0.40 mm) are activated using O_2_ plasma (p_i_ = 0.3 mbar, 100 W, 1 min) and immediately transferred into dry (4 Å MS) toluene (50 mL) under argon. 3-(trimethoxysilyl) propyl methacrylate (1 mL) is added, and the reaction mixture heated to 50°C for 24 h. The slides are then cooled to room temperature and washed by sonication with 3 × 10 mL fresh toluene. The slides are then dried under vacuum in a silicone-free vacuum oven (50°C) for 24 h.

### Molding of TMPMP-co-TEGDA Substrate

TEGDA (337 µL) and TMPMP (163 µL) are added together under argon with DMPA (16.9 mg). The mixture is then sonicated for 15 min to ensure mixing. Each ChemoTopoChip mold on the silicon wafer is framed on 3 sides with Scotch tape (3M) spacers, and a methacrylate silanized glass slide placed on top of each ChemoTopoChip to be molded; standard glass microscope slides (25 mm × 75 mm × 1.0 mm) are placed on top as weights to hold the silanized slides in place. The TMPMP/ TEGDA reaction mixture is transferred into an argon glove box (< 2000 ppm O_2_) along with the silicon mold, and the monomer solution (60 µL) pipetted between the silicon wafer and silanized slides. The rate of pipetting was manually maintained at a similar rate to that of the capillary forces acting upon the solution. When all ChemoTopoChip positions have been pipetted (∼10 min per ChemoTopoChip) they are irradiated with UV light (368 nm, 2 × 15 W bulbs, 10 cm from source) for 10 min. Once complete, the entire molding setup is removed from the glove box and the glass microscope slide weights removed. The silicon wafer is then placed on to a pre-heated (70°C) hot plate; after 10 min, the molded ChemoTopoChips are carefully removed using a scalpel (CAUTION: excessive force and speed will break the thin glass substrate). Once removed, the molded ChemoTopoChips are cleaned by sonication in acetone (10 mL, 10 min) then isopropyl alcohol (10 mL, 10 min). Finally, the ChemoTopoChips are dried under vacuum (0.3 mbar) for 24 hours before functionalization.

### Functionalization of Molded ChemoTopoChip Samples

Monomer solutions are made up as follows: 75% v/v in *N,N*-dimethylformamide (DMF) for oils; 50% w/v in DMF for solids. Next, 0.05% w/v photoinitiator DMPA is added to these solutions before degassing by sonication (10 min). The molded ChemoTopoChip samples are then transferred into an argon glove box (< 2000 ppm O_2_) along with these monomer solutions. A total of 3 µL of monomer solution is then applied to each respective ChemoTopo unit, taking care to evenly cover the entire area required for functionalization. The ChemoTopoChips are then irradiated with UV light (368 nm, 2 × 15 W bulbs, 10 cm from source) for 15 min, before being removed from the argon glove box and sonicated in isopropanol for 10 min. Due to the lower bond dissociation energy of the acrylate π-bond^[1]^ compared with that of the thiol σ-bond,^[2]^ it is expected that these monomers will polymerize to the thiol moieties on the base TMPMP-co-TEGDA substrate after photoinitiation commences. The samples are then placed under vacuum (0.3 mbar) for 7 days before use.

### Time-of-Flight Secondary-Ion Mass Spectrometry (ToF-SIMS) Analysis

ToF-SIMS analysis was carried out using a ToF-SIMS IV (IONTOF GmbH) instrument operated using a 25 kV Bi_3_^+^ primary ion source exhibiting a pulsed target current of ∼1 pA. Samples were scanned at a pixel density of 100 pixels per mm, with 8 shots per pixel over a given area. An ion dose of 2.45 × 10^11^ ions per cm^2^ was applied to each sample area ensuring static conditions were maintained throughout. Both positive and negative secondary ion spectra were collected. Owing to the non-conductive nature of the samples, a low energy (20 eV) electron flood gun was applied to provide charge compensation.

### Atomic Force Microscopy (AFM) Analysis

AFM measurements were conducted using a Bruker Dimension FastScan Bio Icon AFM in Peak Force(tm) (Tapping) mode. Scan areas were 500 × 500 nm and 4 regions of interest (ROIs) were taken. Bruker RTESPA-150 probes were used for all analyses, with all results calibrated to a Bruker polystyrene (2.7 GPa) standard.

### X-Ray Photoelectron Spectroscopy (XPS) Analysis

XPS characterization was carried out using a Kratos AXIS ULTRA DLD. Data was processed using CasaXPS version 2.3.20 rev1.2G

### Mesenchymal Stem Cell Culture

Human immortalized mesenchymal stem cells (hiMSCs) were generated in-house by lentiviral transfection of E6/E7 and hTERT genes as previously described.^[42], [43]^ Cells were cultured in Dulbecco’s modified Eagle’s medium supplemented with 10% (v/v) foetal bovine serum, 100 units/mL penicillin, 100□μg/mL streptomycin and non-essential amino acids. Positive controls were cultured in Human Mesenchymal Stem Cell (hMSC) Osteogenic Differentiation Medium (PT-3002; Lonza). All cells were maintained in a humidified incubator at 37°C and 5% CO_2_ in air. Cells were re-suspended in the appropriate volume of media and seeded on 3 replicate ChemoTopoChips at 1 × 10^5^ hiMSCs/chip (3 independent experiments using cells from 3 different passage numbers).

### hiMSC Immunofluorescence Staining

For alkaline phosphatase (ALP) staining, cells were cultured on the ChemoTopoChips for five days in culture medium (at 37°C, 5% CO_2_ in air) then fixed using 70% (v/v) ethanol, permeabilized with 0.1% (v/v) Triton X-100 and incubated with a blocking solution of 3% (v/v) goat serum in 1% (v/v) BSA/PBS. Staining was carried out using human ALP antibody (Dilution 1:50; sc137213, Santa Cruz Biotech) and counterstained for α-tubulin (2 µg/mL; PA120988, Invitrogen) for 3 hours at room temperature. After washing, slides were incubated with the appropriate secondary antibodies in the green and red channels at room temperature (1:100 dilution). Nuclei were stained with NucBlue Fixed Cell ReadyProbes(tm) (Invitrogen).

### Monocyte Isolation and Culture

Buffy coats were obtained from the National Blood Service after obtaining written informed consent and approval from the ethics committee. Monocytes were isolated from□peripheral blood mononuclear cells (PBMCs). A MACS magnetic cell separation system (CD14□MicroBeads□positive selection with LS columns, Miltenyi Biotec) was used for the isolation as previously described. Isolated monocytes were prepared in RPMI-1640 medium containing 10%□foetal bovine serum (FBS), 100□µg/ml□streptomycin, 2mM L-glutamine and 100 U/ml penicillin (Sigma-Aldrich). For assessment of cell attachment and phenotype characterization, cells were re-suspended in the appropriate volume of media and seeded on the ChemoTopoChips at 2 × 10^6^ monocytes/chip and incubated at 37°C, 5% CO_2_□in a humidified incubator for 9 days.□

### Macrophage Immunofluorescent Staining

On day 9 all adherent cells cultured on ChemoTopoChips were fixed in 4% paraformaldehyde (BioRad) in PBS, then blocked with 3% BSA (Sigma-Aldrich) and 1% Glycine (Fisher Scientific) in PBS. Subsequently, another blocking step was carried out using 5% goat serum (Sigma) in PBS. Adherent cells were stained with□2 µg/mL□anti-human TNFα (IgG1) mAb (Abcam), and with□1 µg/ml□anti-human IL-10 (IgG1) mAb (Abcam) followed by 1 h incubation at room temperature. After washing, cells were stained with□8 µg/ml□Rhodamine-x goat anti-mouse IgG (H+L) secondary Ab (Invitrogen), and□8 µg/ml□Alexa flour-647 goat anti-rabbit IgG (H+L) secondary antibody (Invitrogen) for another hour at room temperature. All samples were counterstained with 250 ng/ml DAPI (4’,6-Diamidino-2-Phenylindole) (Invitrogen) at room temperature.

### ChemoTopoChip Imaging

Imaging of all fixed and stained ChemoTopoChip samples was carried out using a widefield deconvolution-TIRF3 system (Zeiss, custom setup). Imaging was carried out in wide field mode using a 20×/0.5 NA air objective in the bright field and fluorescence channels with the excitation at 358 nm, 488nm and 561 nm. The software used to capture was Zeiss Zen Blue, by using the “Sample Carrier Designer” wizard/module to manually create and calibrate the position list which was used to scan all the positions in the chip setup.

### CellProfiler Analysis

A custom CellProfiler pipeline was created to correct for uneven background illumination in each image, then each image cropped to within the Topo unit 30 µm wall. Nuclei were detected using an adaptive per-object algorithm in the blue channel images, followed by propagation from these primary detected objects to detect cell cytoskeleton and ALP staining (hiMSCs) or TNFα and IL-10 (human macrophages) in the green and red channel images. Intensity of detected objects was measured and exported, and images containing overlaid outlines of detected objects also saved to ensure correct operation of the pipeline.

### Statistical Analysis

Statistical anlysis and graphical plots were carried out in R version 3.6.1 using RStudio version 1.2.1335 as integrated development environment (IDE). For an exploratory method combinations having p-value < 0.05 were highlighted from a two independent sample equal variance t-test. Note that the sample sizes are small here and this does not account for multiple tests. The values were normalized to the base polymer region on each slide. Heatmaps were plotted using the heatmap.2 function from the gplots package version 3.1.0.2 in combination with the RColorBrewer package version 1.1-2. Clustering and dendrograms for heatmaps were produced using the complete linkage method^[3]^ with Euclidean distance measure.

Ranked scatter plots and box plots were carried out using base functions in R and the ggplot2 package version 3.2.1.

### Synergy Ratio Determination

Assessment of the interactions between binary factors (chemistry and topography) is readily performed using a synergy ratio (*SR*). Taking the response of factor *x*_1_ alone (*y*_1_), the response of factor *x*_2_ alone (*y*_2_) and the response of the factors combined *x*_12_ (*y*_12_), *SR* can be calculated as shown in Equation S1:

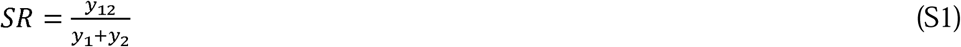

For a synergistic combination, *SR* > 1 as the ratio is then greater than the sum of the theoretical maximums of the individual response factor comparators; for a counteractive combination, SR < 0.5 as the ratio is then less than the theoretical maximum of one individual response factor alone (i.e. 0 contribution from the second individual response factor). In analysis of the ChemoTopoChip data, unfunctionalized TMPMP-co-TEGDA molded topographies and flat area chemistries were used as the individual factors *x*_1_ and *x*_2_ to compare with the hit topography-material combinations *x*_12_.

### Random Forest Machine Learning

The raw dataset consisted of three technical repeats for each surface variable (topography, chemistry) within a chip, which were further replicated across multiple batches (biological repeats). Data set from repeats in a chip have been normalized against their correspondent flat values. Subsequently, replicate average values were calculated. The average between batches was then determined as the dependent variable for the predictive models. Macrophage polarization and ALP intensity predictive models were generated.

The various topographies were encoded using descriptors generated by CellProfiler^[4]^ that relate directly to particular primitives in the topographical units. For chemistries, 1-hot descriptors (binary variables indicating the presence or absence of a chemistry in any given combination) were used.

The SHapley Additive exPlanation (SHAP) method was used for feature selection to eliminate uninformative and less informative descriptors and less relevant chemistries. SHAP was implemented using the SHAP package in Python 3.7. Regression models were generated using the random forest approach with the scikit-learn package in Python 3.7. The default parameters from version 0.22 were adopted for the random forest models. That is, 100 estimators were considered using gini as the function to measure the quality of the data instances split. And no limit for the maximum depth of the trees was defined. 70% of the data instances were employed for model training and 30% for testing. The performance of the predictive models and the topographical descriptors that contributed most strongly to the attachment and polarization are shown in **Figure 3**. The figure presents the results of the regression models as well as the features selected. The features are ordered from top to bottom based on their average impact on the model output magnitude

**Figure S1.**
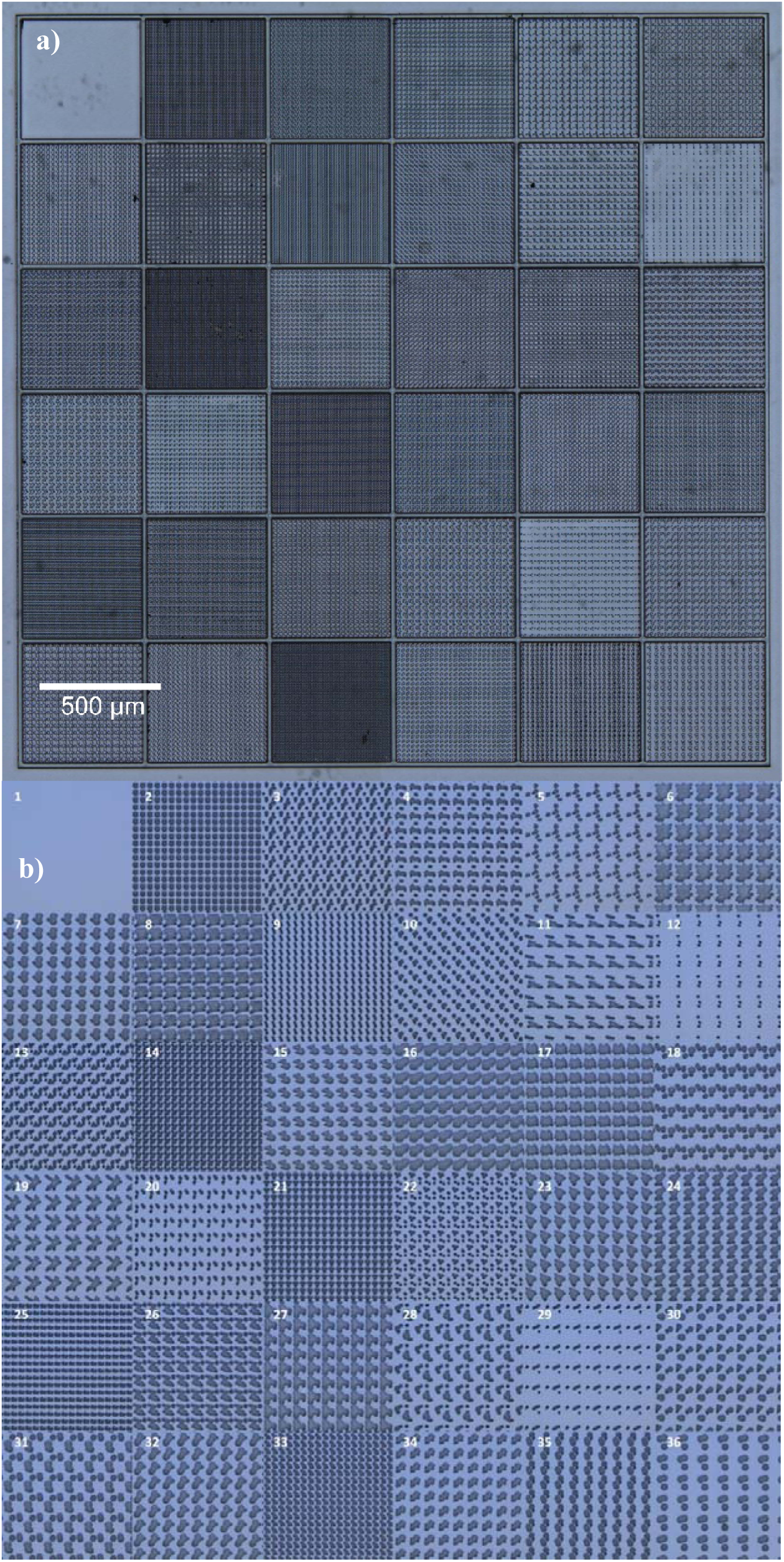
a) Optical profiler image of ChemoTopoChip b) Images of topography shapes.

**Figure S2.**
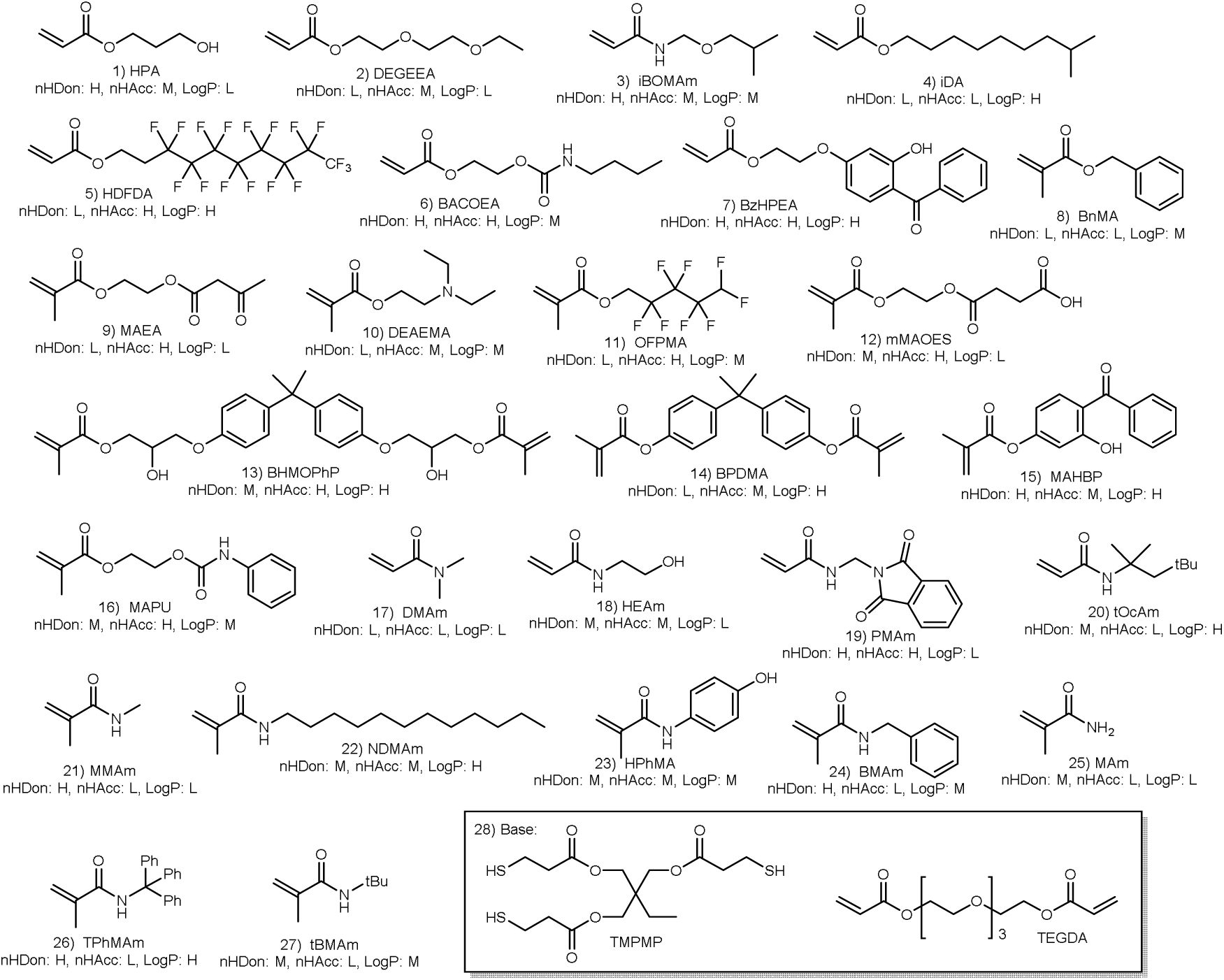
Chemistries Used in the ChemoTopoChip: nHDon, nHAcc and LogP refer to number of H-bond donors, number of H-bond acceptors and LogP (octanol/water partition coefficient) classified as high (H), medium (M) or low (L)

**Figure S3.**
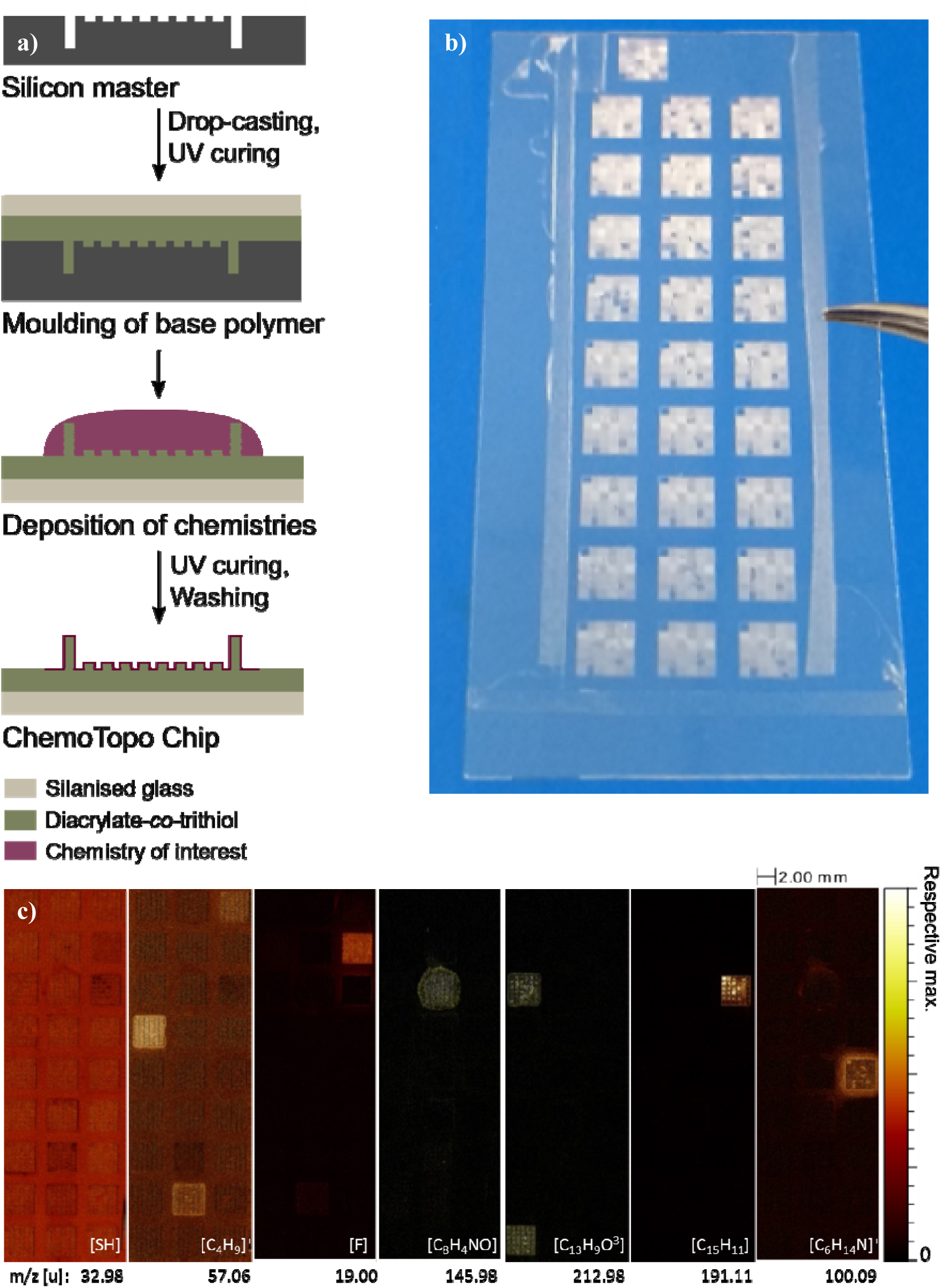
ChemoTopoChip Manufacture. **a)** ChemoTopoChip production process; **b)** Photographed ChemoTopoChip; **c)** ToF-SIMS images of functionalized surface showing the distribution of the thiol ion from the base and of 6 ions unique to specific functionalization chemistries.

**Figure S4.**
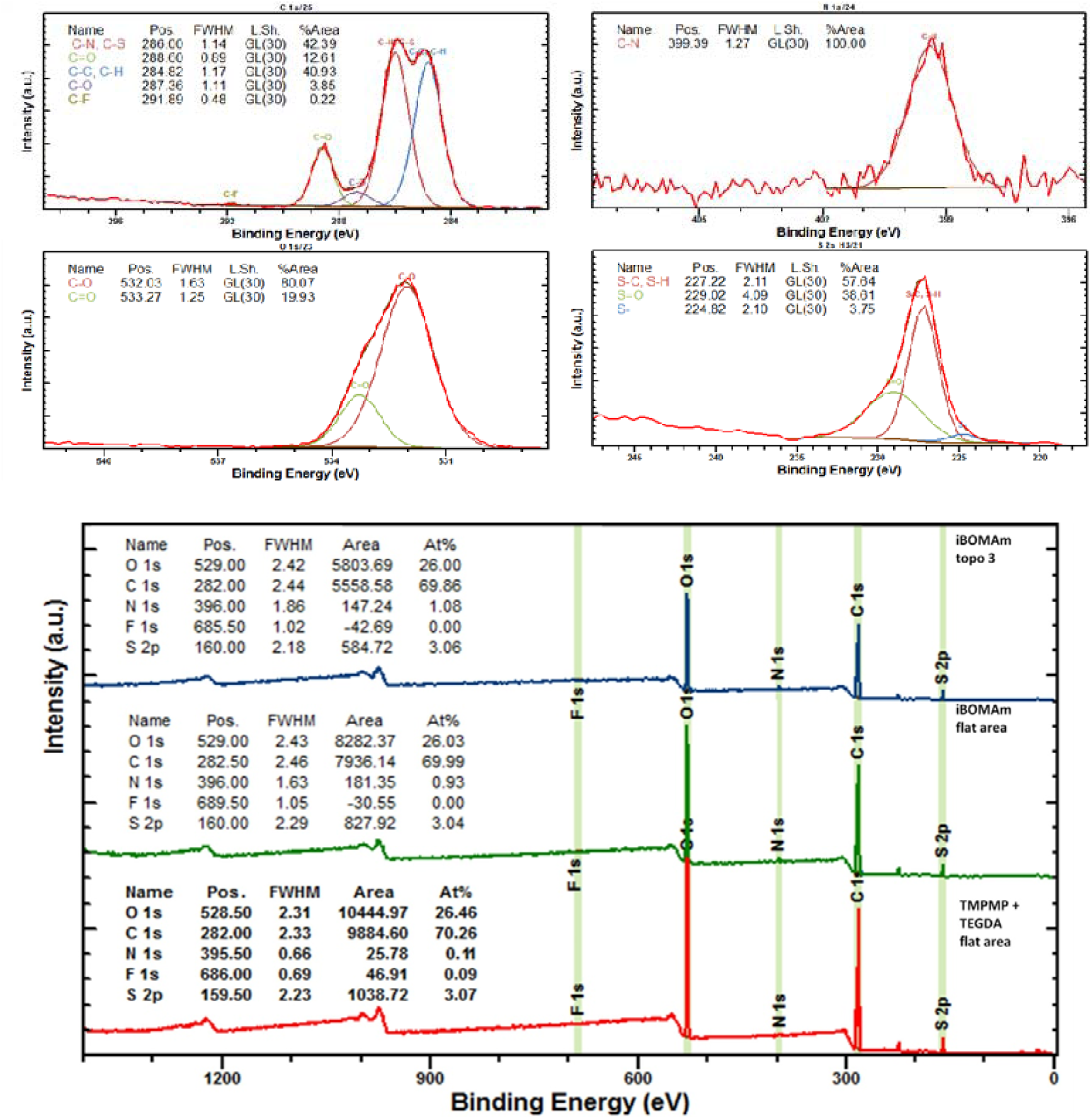
XPS of Base TMPMP-co-TEGDA and Example iBOMAm Functionalized Area.

**Figure S5.**
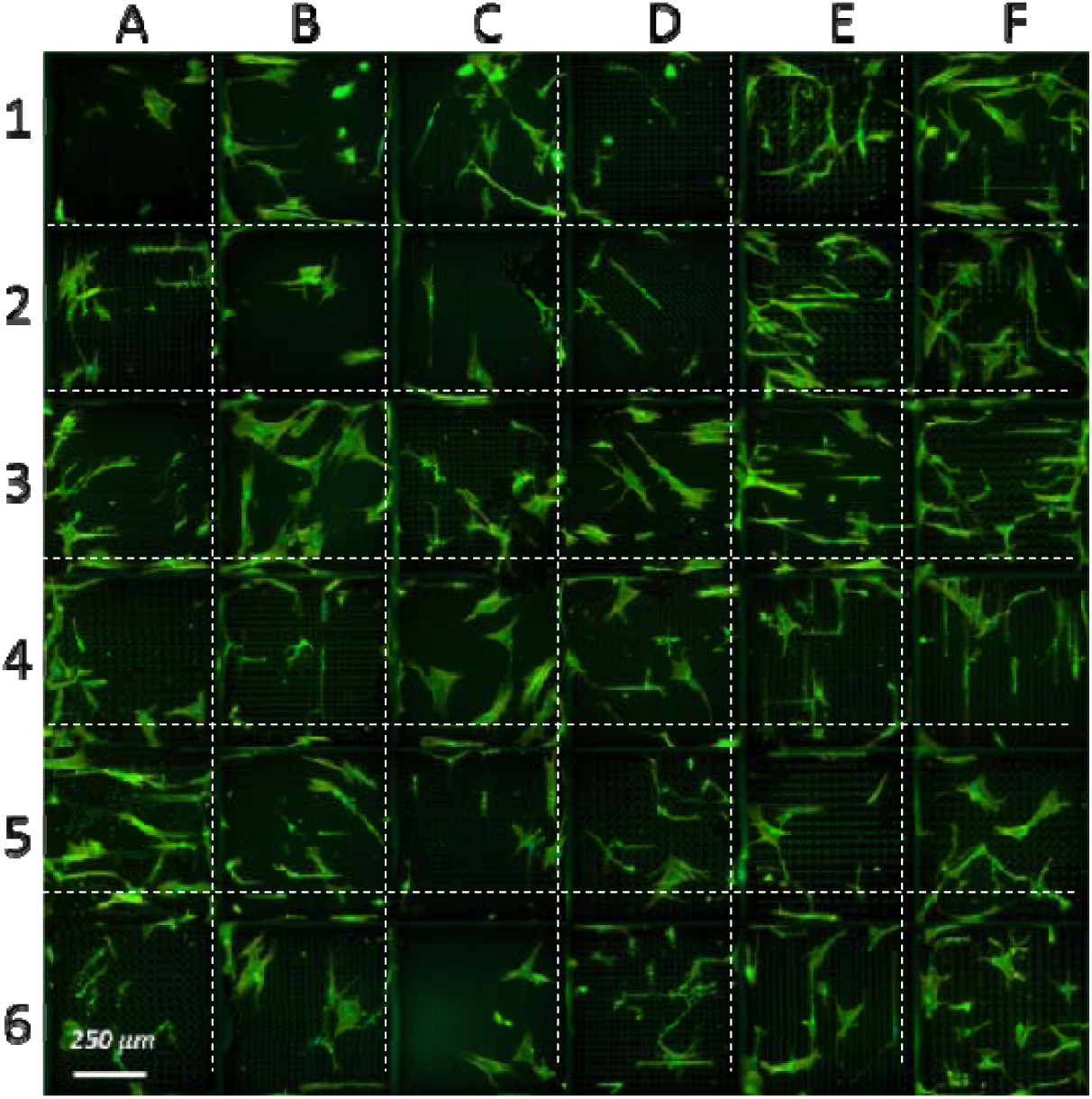
TMPMP-co-TEGDA ChemoTopo unit, stained with α-tubulin.

**Figure S6.**
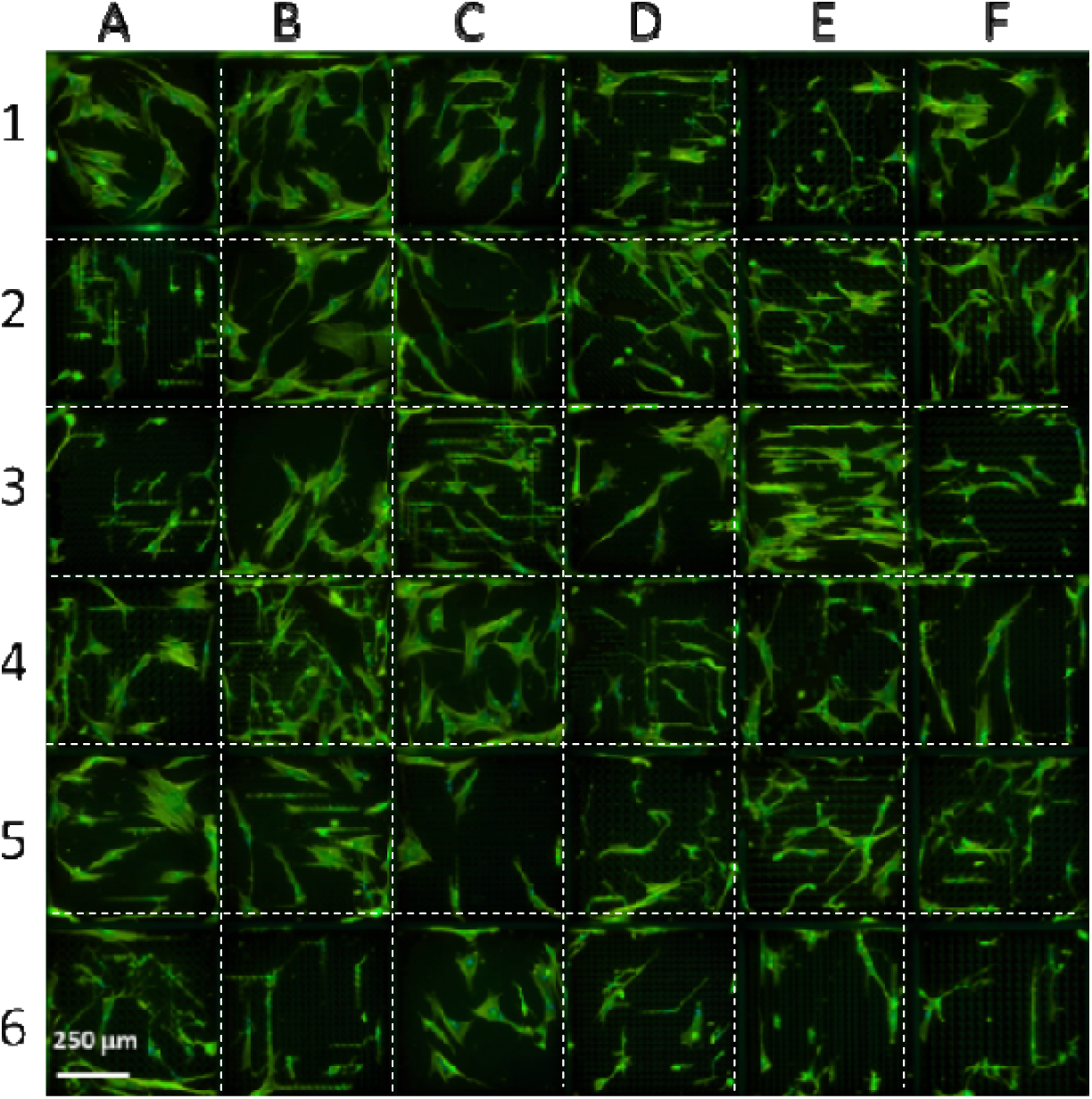
iBOMAm ChemoTopo unit, stained with α-tubulin.

**Figure S7.**
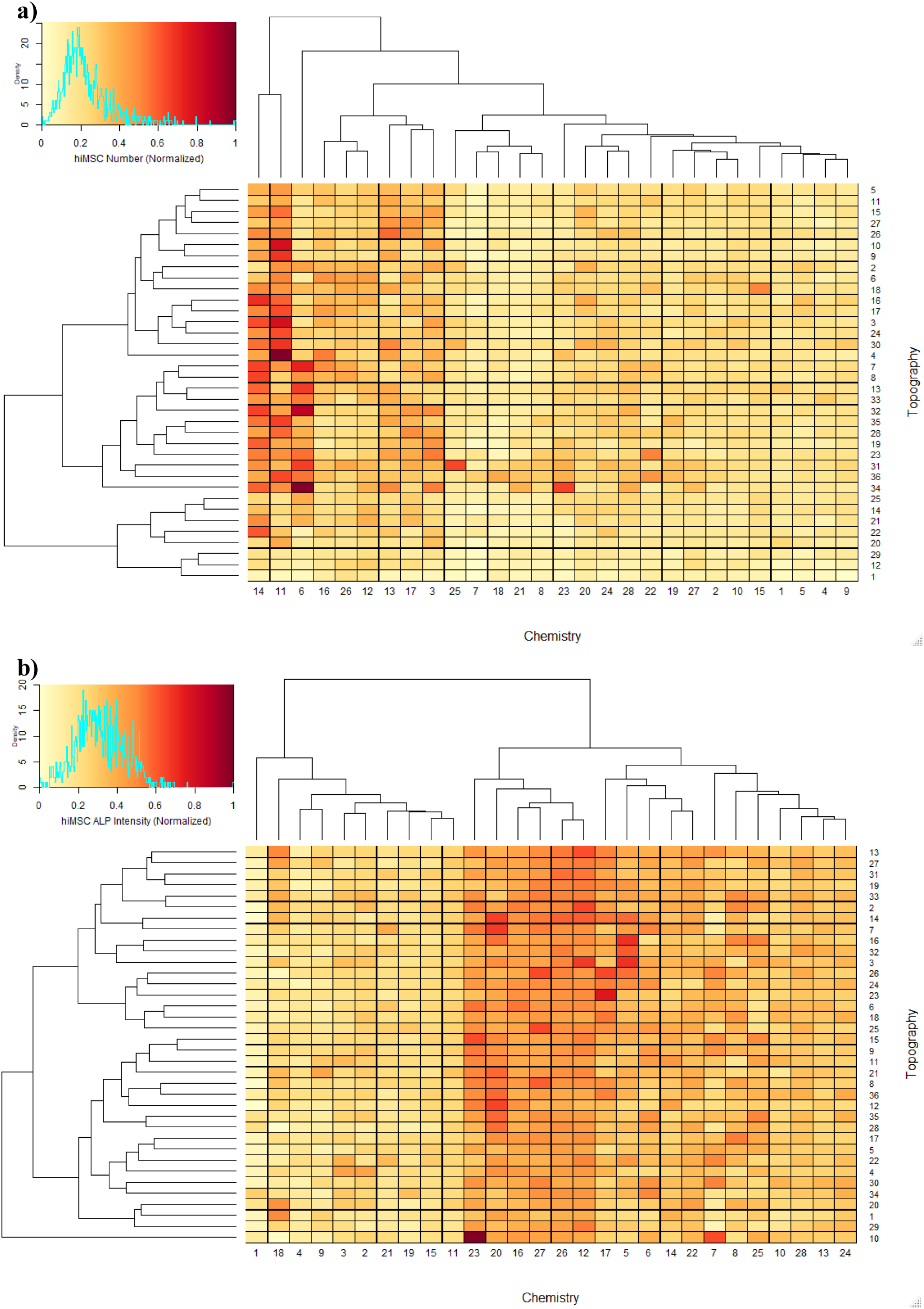
Normalized hiMSC a) Number and b) Mean Integrated ALP Expression Across ChemoTopoChip.

**Figure S8.**
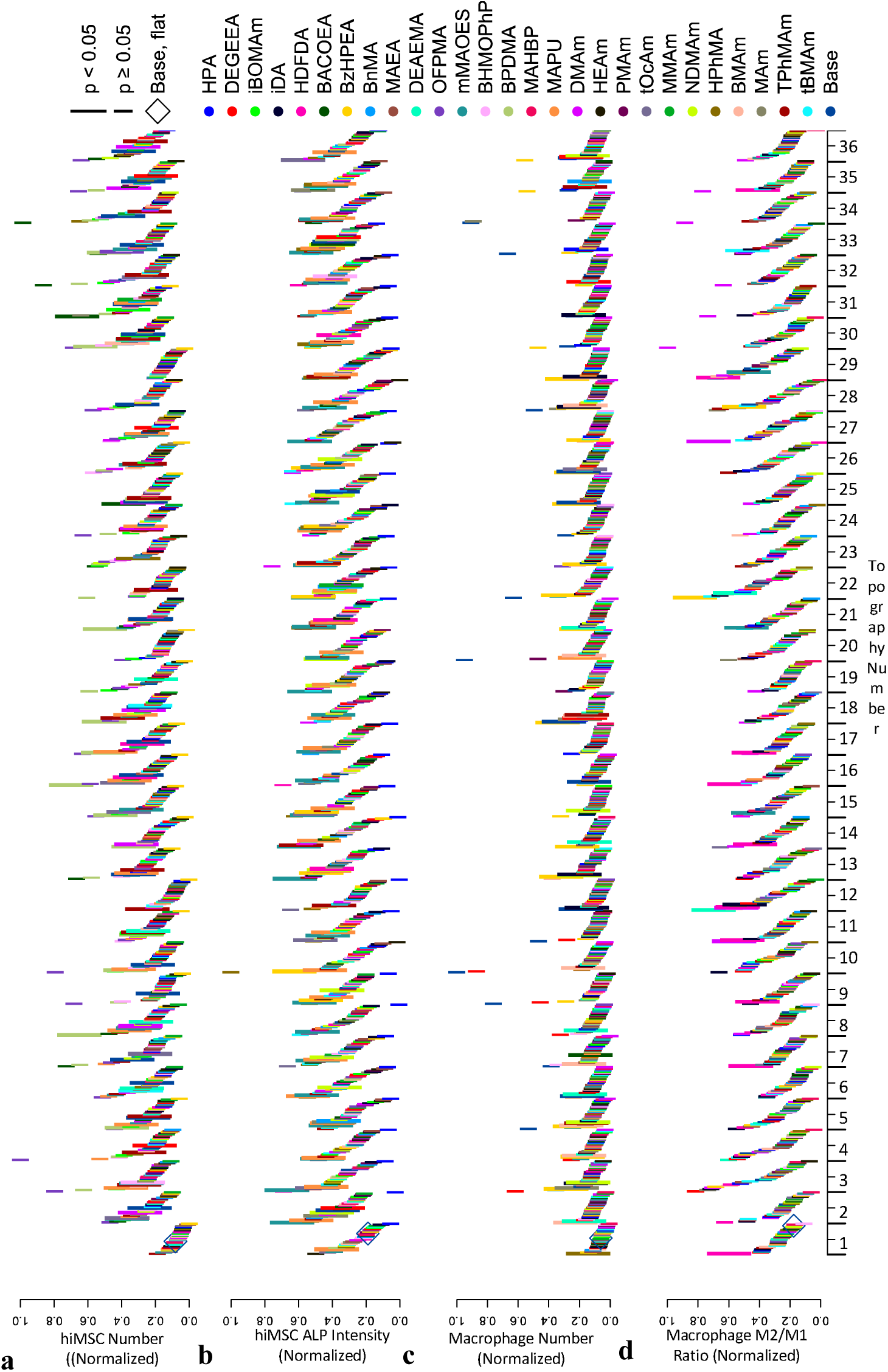
Rank ordered a) hiMSC cell number (N = 3, n = 3) b) hiMSC ALP intensities (N = 3, n = 3) c) human macrophage cell number (N = 2, n = 3) and d) human macrophage M2/M1 ratio (N = 2, n = 3) normalized to flat TMPMP-co-TEGDA area on each ChemoTopoChip.

**Figure S9.**
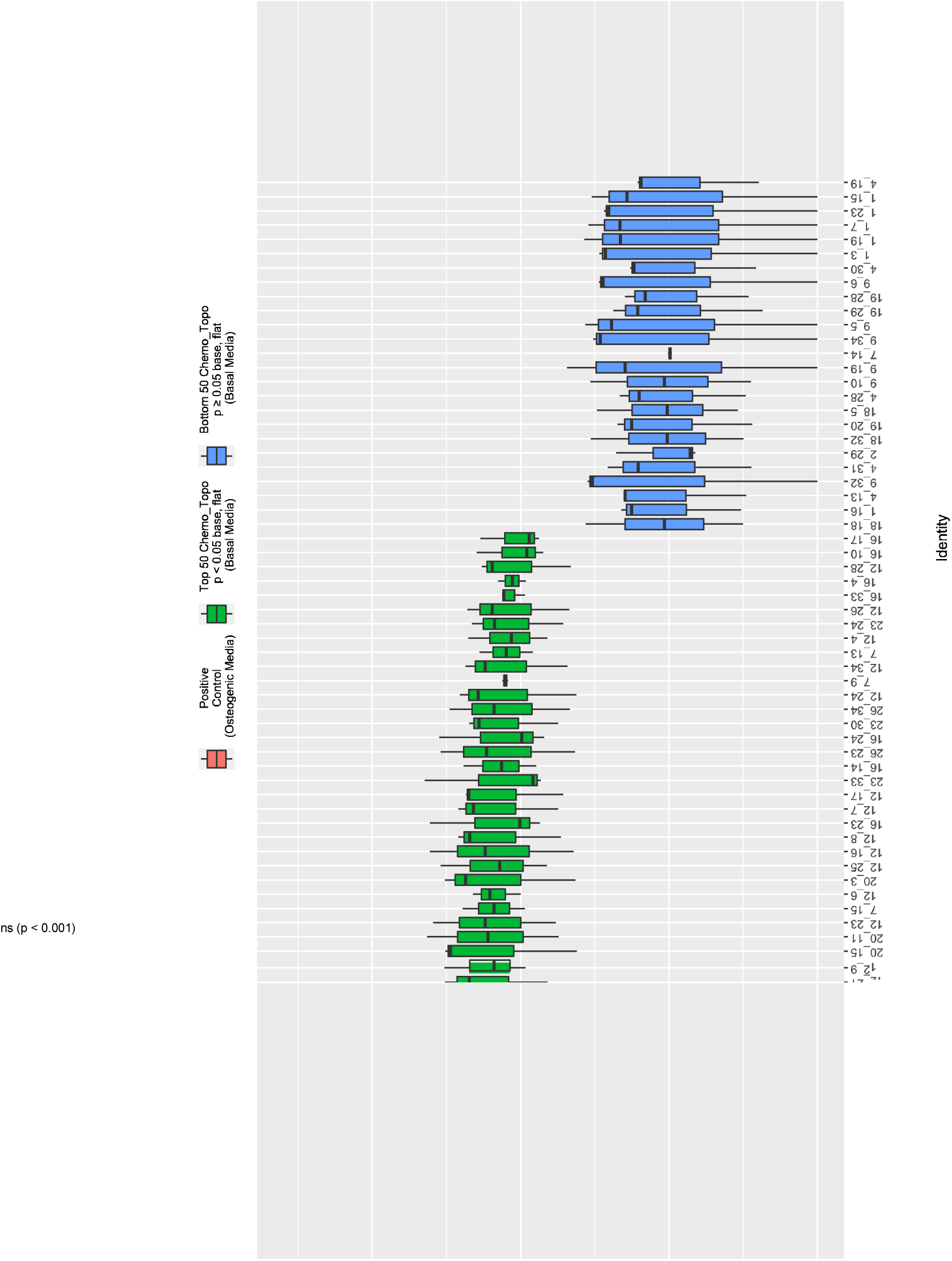
Raw ALP Intensities of Top 50 ChemoTopo Combinations (p < 0.05) and Bottom 50 ChemoTopo Combinations (p ≥ 0.05), Compared to Flat Base TPMP-co-TEGDA Region, and Positive Control.

**Figure S10.**
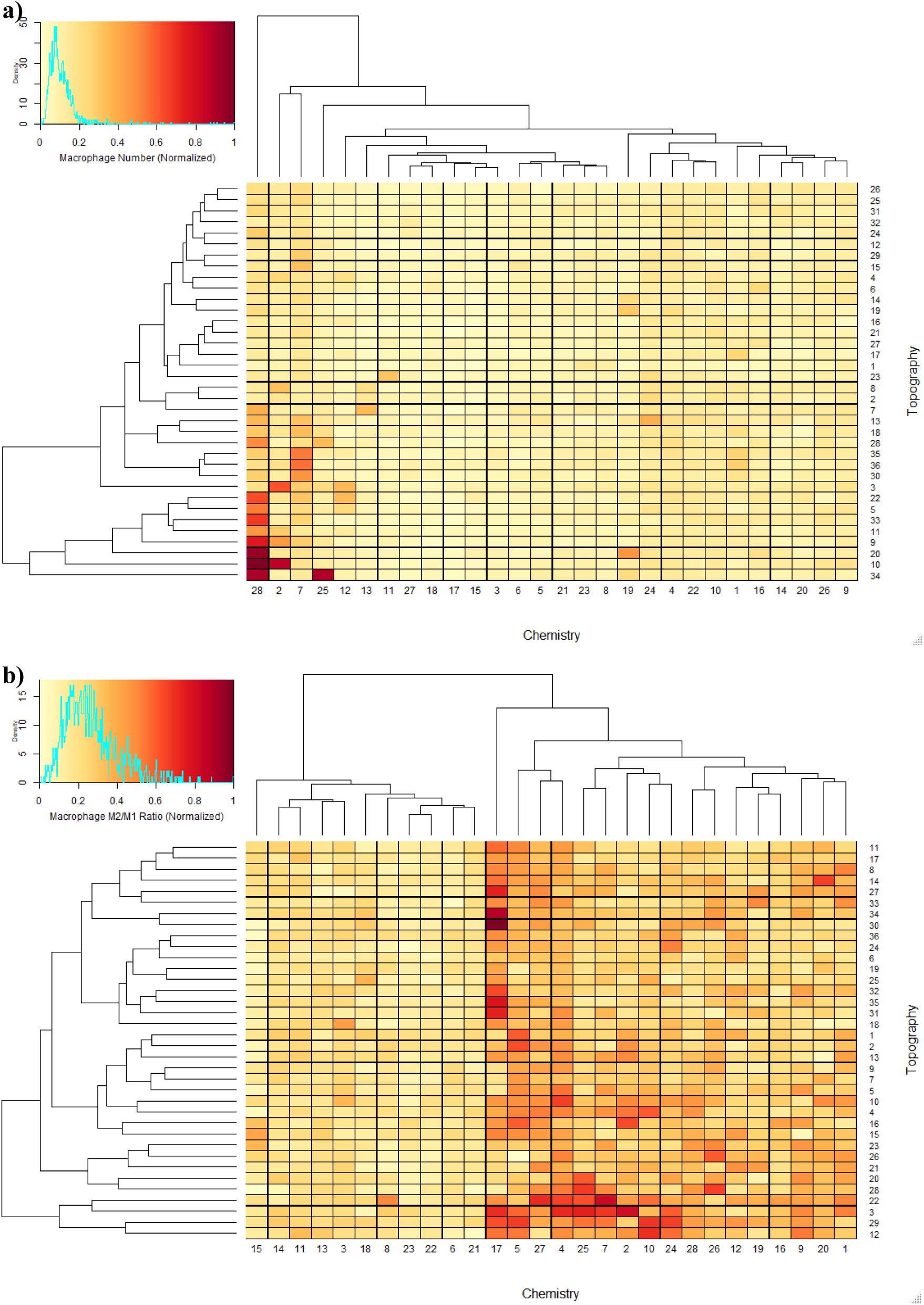
Normalized Macrophage a) Number and b) M2/M1 Ratio Across ChemoTopoChip.

**Figure S11.**
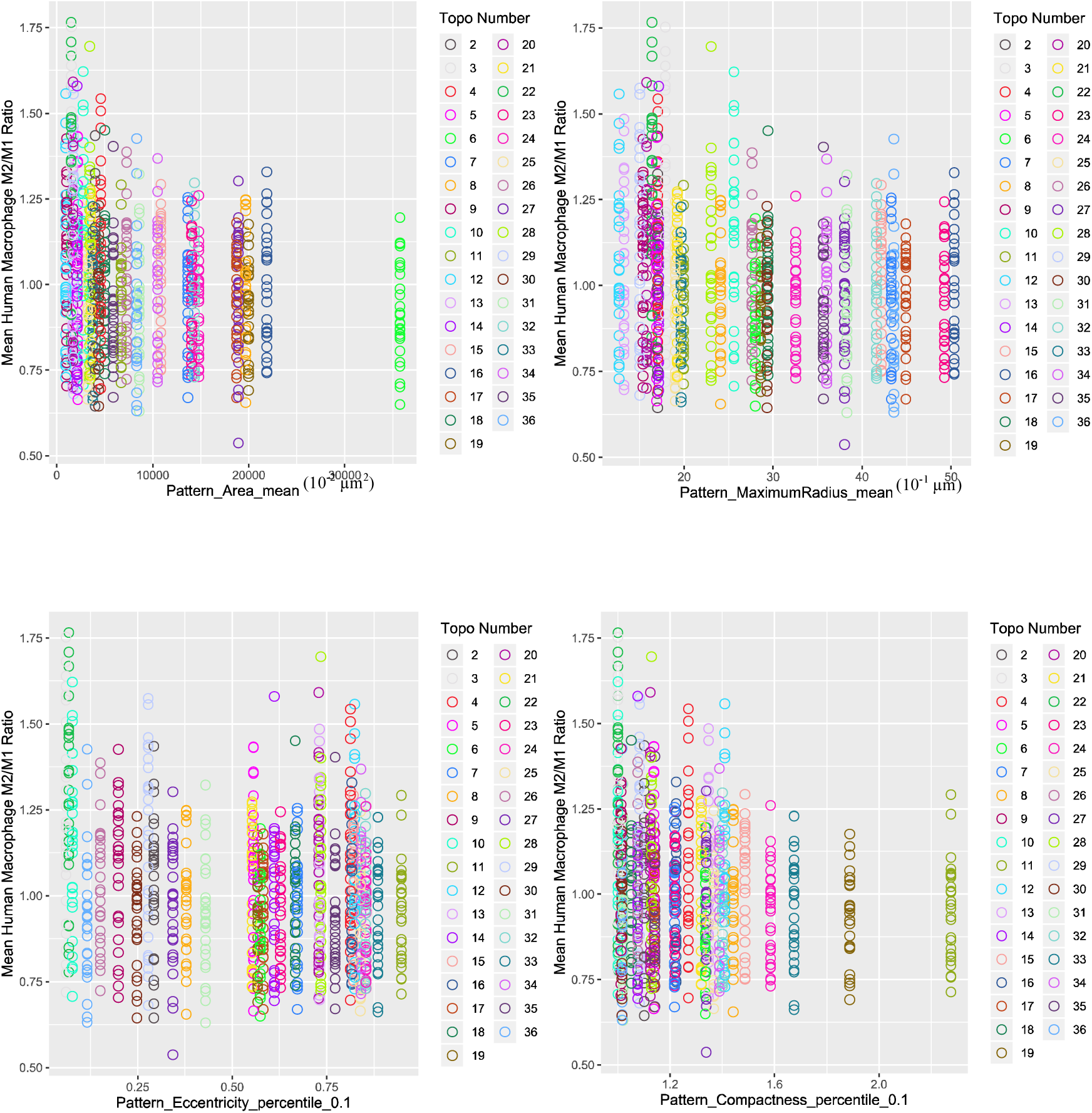
Human Macrophage M2/M1 Ratio Topographical Descriptor Correlation Plots.

**Figure S12.**
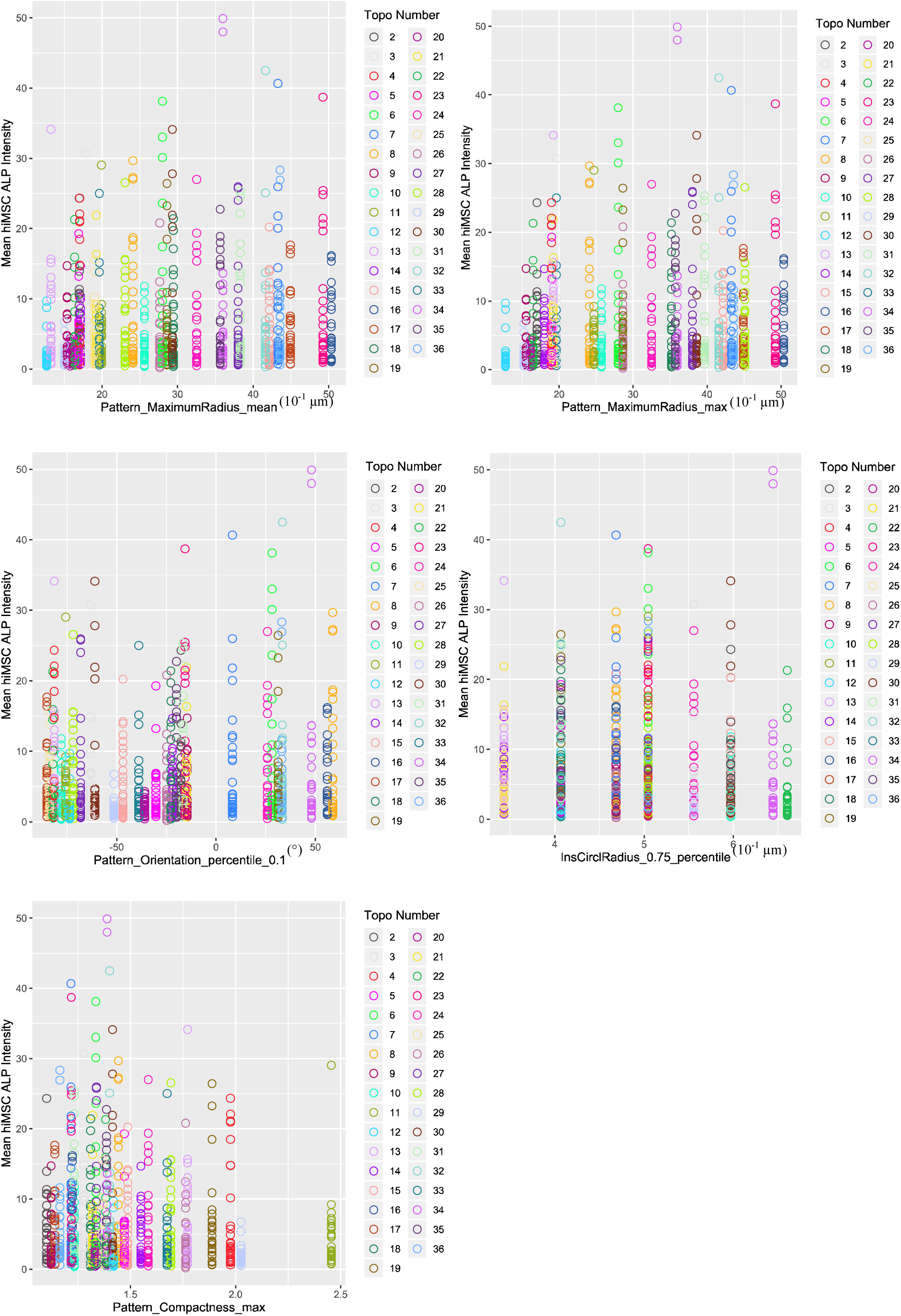
hiMSC ALP Intensity Topographical Descriptor Correlation Plots.

**Table S1.**
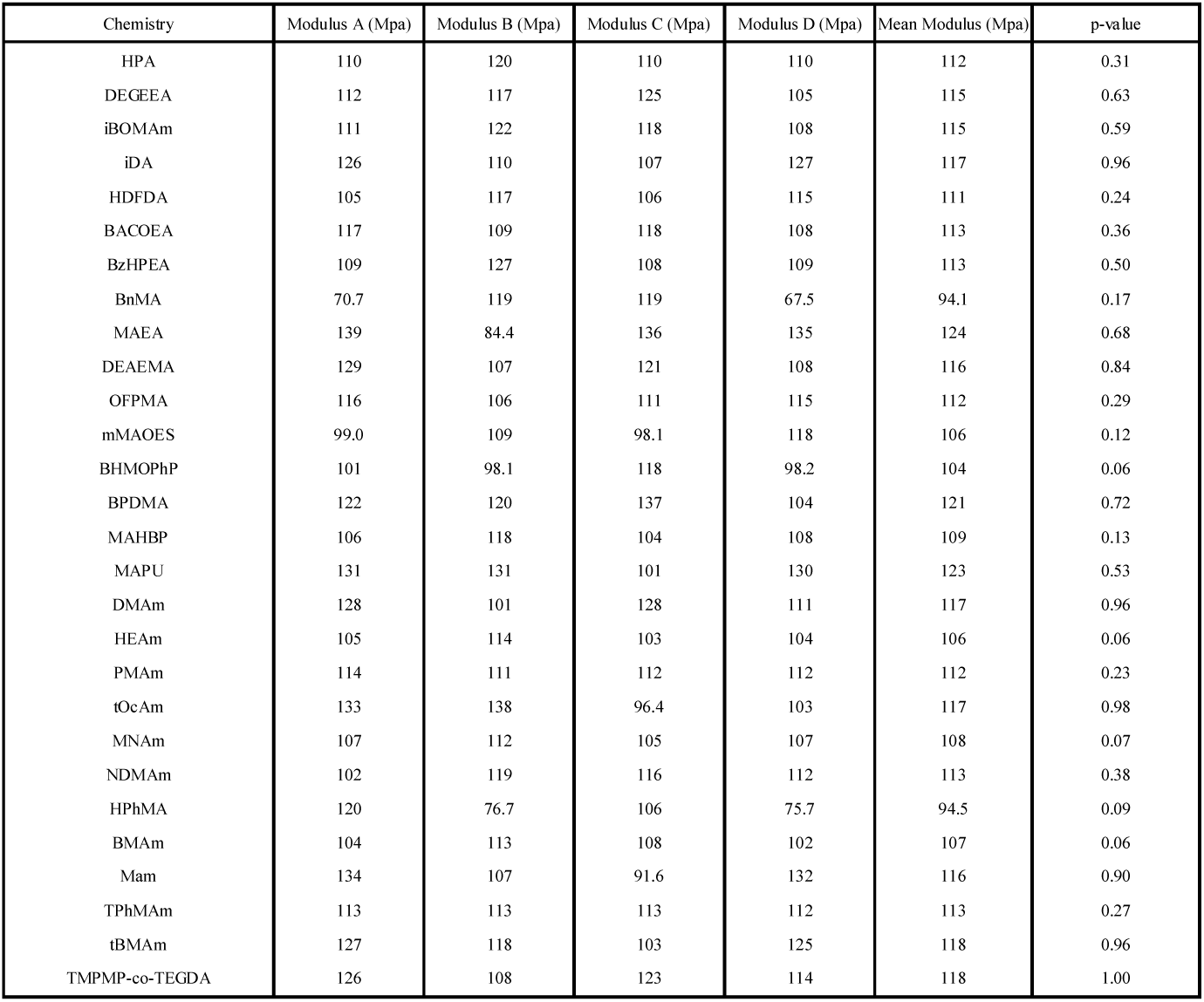
AFM Modulus data taken from 4 ROIs.

**Table S2.**
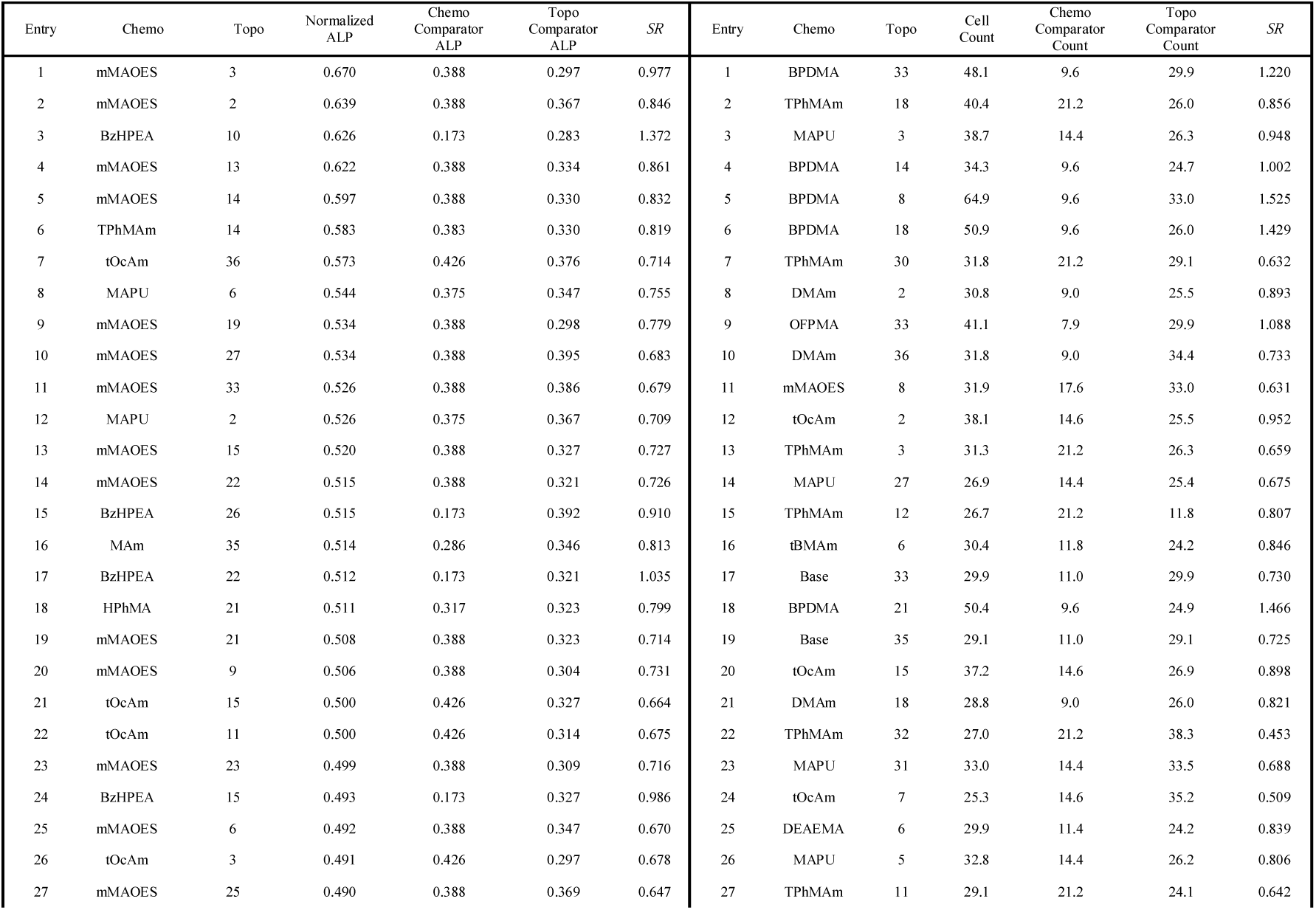
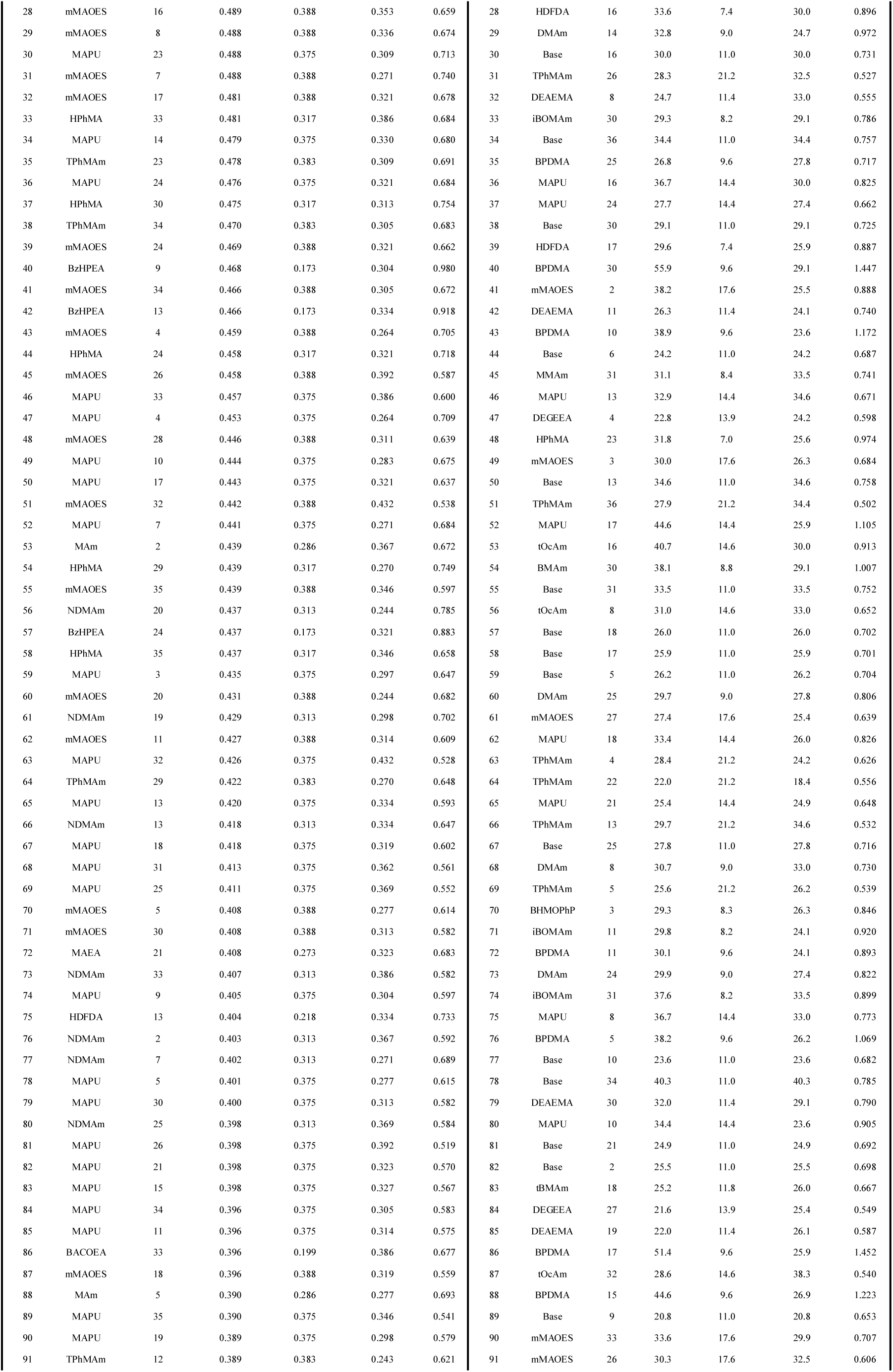
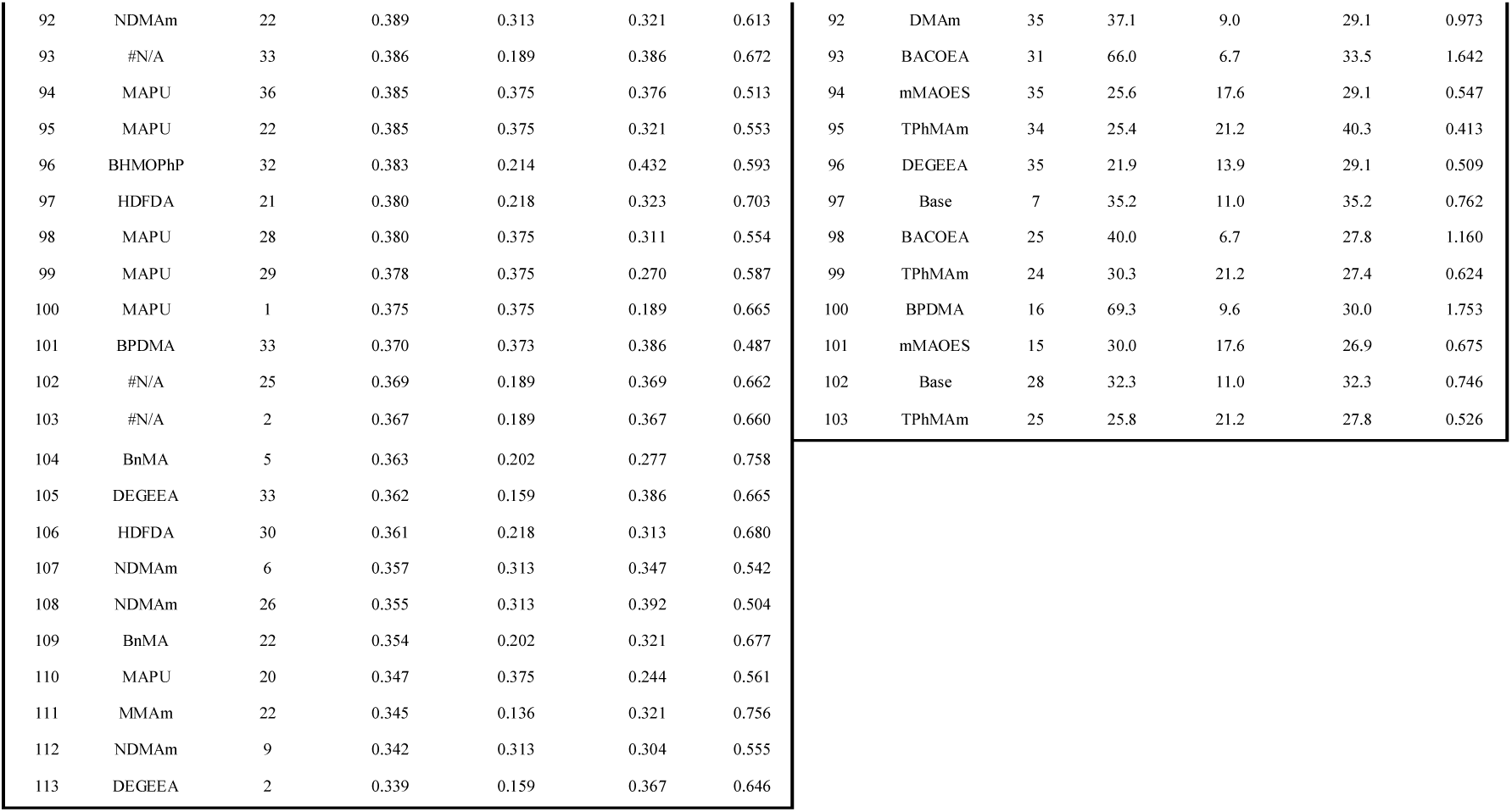
Statistically Signification hiMSC Combinations.

**Table S3.**
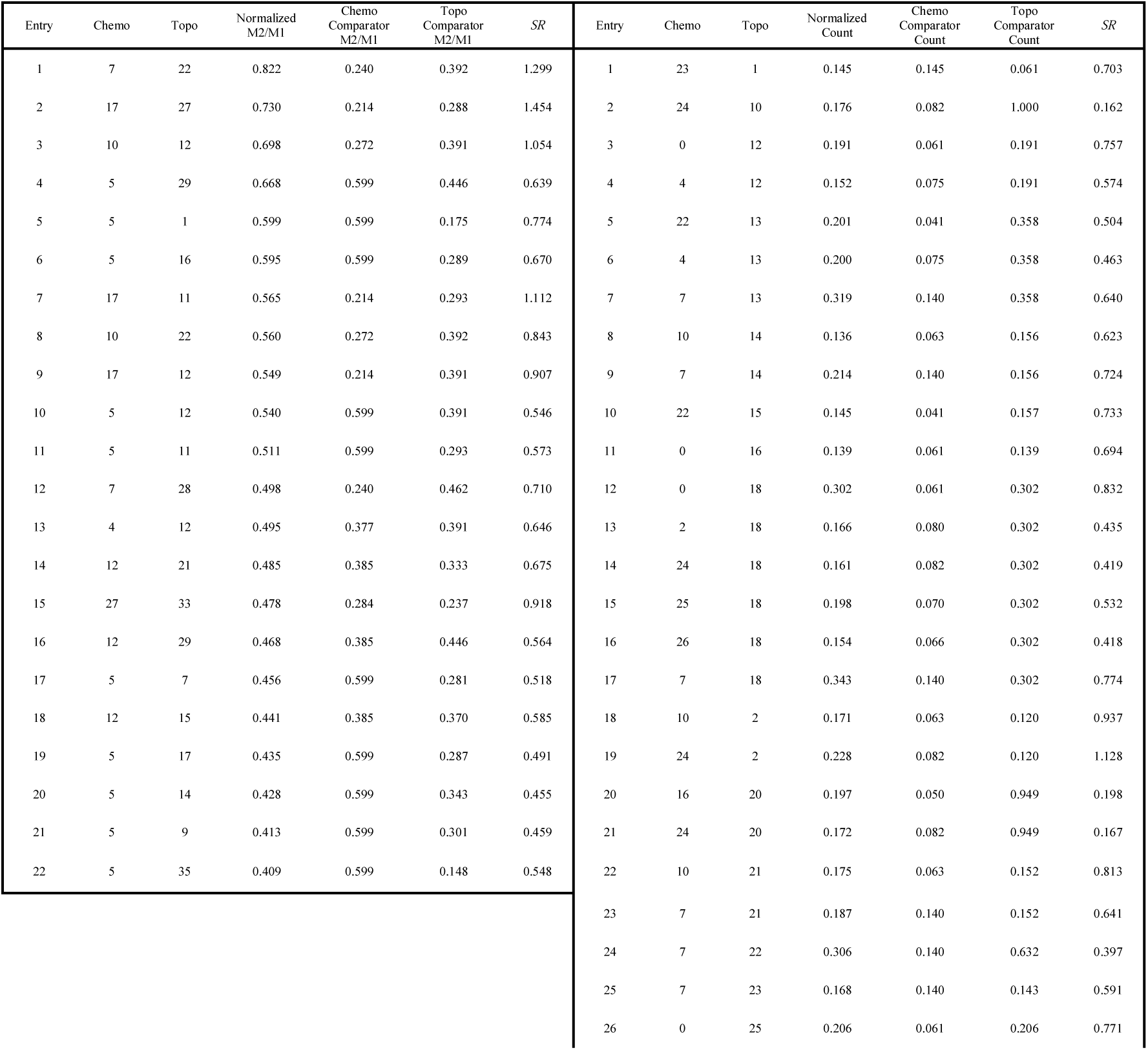
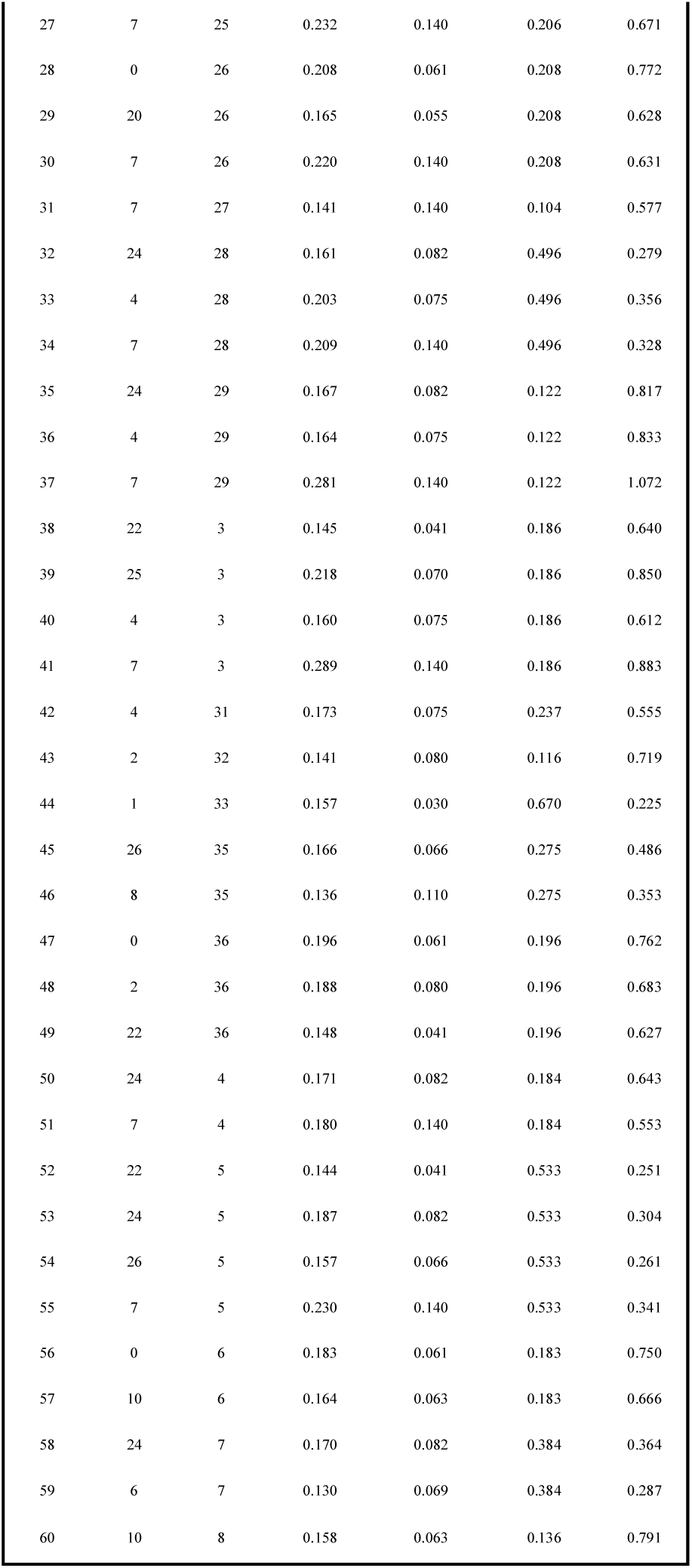
Statistically Significant Macrophage Combinations.

| Entry | Chemo | Topo | Normalized M2/M1 | Chemo Comparator M2/M1 | Topo Comparator M2/M1 | SR | Entry | Chemo | Topo | Normalized Count | Chemo Comparator Count | Topo Comparator Count | SR |
| --- | --- | --- | --- | --- | --- | --- | --- | --- | --- | --- | --- | --- | --- |
| 1 | 7 | 22 | 0.822 | 0.240 | 0.392 | 1.299 | 1 | 23 | 1 | 0.145 | 0.145 | 0.061 | 0.703 |
| 2 | 17 | 27 | 0.730 | 0.214 | 0.288 | 1.454 | 2 | 24 | 10 | 0.176 | 0.082 | 1.000 | 0.162 |
| 3 | 10 | 12 | 0.698 | 0.272 | 0.391 | 1.054 | 3 | 0 | 12 | 0.191 | 0.061 | 0.191 | 0.757 |
| 4 | 5 | 29 | 0.668 | 0.599 | 0.446 | 0.639 | 4 | 4 | 12 | 0.152 | 0.075 | 0.191 | 0.574 |
| 5 | 5 | 1 | 0.599 | 0.599 | 0.175 | 0.774 | 5 | 22 | 13 | 0.201 | 0.041 | 0.358 | 0.504 |
| 6 | 5 | 16 | 0.595 | 0.599 | 0.289 | 0.670 | 6 | 4 | 13 | 0.200 | 0.075 | 0.358 | 0.463 |
| 7 | 17 | 11 | 0.565 | 0.214 | 0.293 | 1.112 | 7 | 7 | 13 | 0.319 | 0.140 | 0.358 | 0.640 |
| 8 | 10 | 22 | 0.560 | 0.272 | 0.392 | 0.843 | 8 | 10 | 14 | 0.136 | 0.063 | 0.156 | 0.623 |
| 9 | 17 | 12 | 0.549 | 0.214 | 0.391 | 0.907 | 9 | 7 | 14 | 0.214 | 0.140 | 0.156 | 0.724 |
| 10 | 5 | 12 | 0.540 | 0.599 | 0.391 | 0.546 | 10 | 22 | 15 | 0.145 | 0.041 | 0.157 | 0.733 |
| 11 | 5 | 11 | 0.511 | 0.599 | 0.293 | 0.573 | 11 | 0 | 16 | 0.139 | 0.061 | 0.139 | 0.694 |
| 12 | 7 | 28 | 0.498 | 0.240 | 0.462 | 0.710 | 12 | 0 | 18 | 0.302 | 0.061 | 0.302 | 0.832 |
| 13 | 4 | 12 | 0.495 | 0.377 | 0.391 | 0.646 | 13 | 2 | 18 | 0.166 | 0.080 | 0.302 | 0.435 |
| 14 | 12 | 21 | 0.485 | 0.385 | 0.333 | 0.675 | 14 | 24 | 18 | 0.161 | 0.082 | 0.302 | 0.419 |
| 15 | 27 | 33 | 0.478 | 0.284 | 0.237 | 0.918 | 15 | 25 | 18 | 0.198 | 0.070 | 0.302 | 0.532 |
| 16 | 12 | 29 | 0.468 | 0.385 | 0.446 | 0.564 | 16 | 26 | 18 | 0.154 | 0.066 | 0.302 | 0.418 |
| 17 | 5 | 7 | 0.456 | 0.599 | 0.281 | 0.518 | 17 | 7 | 18 | 0.343 | 0.140 | 0.302 | 0.774 |
| 18 | 12 | 15 | 0.441 | 0.385 | 0.370 | 0.585 | 18 | 10 | 2 | 0.171 | 0.063 | 0.120 | 0.937 |
| 19 | 5 | 17 | 0.435 | 0.599 | 0.287 | 0.491 | 19 | 24 | 2 | 0.228 | 0.082 | 0.120 | 1.128 |
| 20 | 5 | 14 | 0.428 | 0.599 | 0.343 | 0.455 | 20 | 16 | 20 | 0.197 | 0.050 | 0.949 | 0.198 |
| 21 | 5 | 9 | 0.413 | 0.599 | 0.301 | 0.459 | 21 | 24 | 20 | 0.172 | 0.082 | 0.949 | 0.167 |
| 22 | 5 | 35 | 0.409 | 0.599 | 0.148 | 0.548 | 22 | 10 | 21 | 0.175 | 0.063 | 0.152 | 0.813 |
|  |  |  |  |  |  |  | 23 | 7 | 21 | 0.187 | 0.140 | 0.152 | 0.641 |
|  |  |  |  |  |  |  | 24 | 7 | 22 | 0.306 | 0.140 | 0.632 | 0.397 |
|  |  |  |  |  |  |  | 25 | 7 | 23 | 0.168 | 0.140 | 0.143 | 0.591 |
|  |  |  |  |  |  |  | 26 | 0 | 25 | 0.206 | 0.061 | 0.206 | 0.771 |

|  |  |  |  |  |  |  |
| --- | --- | --- | --- | --- | --- | --- |
| 27 | 7 | 25 | 0.232 | 0.140 | 0.206 | 0.671 |
| 28 | 0 | 26 | 0.208 | 0.061 | 0.208 | 0.772 |
| 29 | 20 | 26 | 0.165 | 0.055 | 0.208 | 0.628 |
| 30 | 7 | 26 | 0.220 | 0.140 | 0.208 | 0.631 |
| 31 | 7 | 27 | 0.141 | 0.140 | 0.104 | 0.577 |
| 32 | 24 | 28 | 0.161 | 0.082 | 0.496 | 0.279 |
| 33 | 4 | 28 | 0.203 | 0.075 | 0.496 | 0.356 |
| 34 | 7 | 28 | 0.209 | 0.140 | 0.496 | 0.328 |
| 35 | 24 | 29 | 0.167 | 0.082 | 0.122 | 0.817 |
| 36 | 4 | 29 | 0.164 | 0.075 | 0.122 | 0.833 |
| 37 | 7 | 29 | 0.281 | 0.140 | 0.122 | 1.072 |
| 38 | 22 | 3 | 0.145 | 0.041 | 0.186 | 0.640 |
| 39 | 25 | 3 | 0.218 | 0.070 | 0.186 | 0.850 |
| 40 | 4 | 3 | 0.160 | 0.075 | 0.186 | 0.612 |
| 41 | 7 | 3 | 0.289 | 0.140 | 0.186 | 0.883 |
| 42 | 4 | 31 | 0.173 | 0.075 | 0.237 | 0.555 |
| 43 | 2 | 32 | 0.141 | 0.080 | 0.116 | 0.719 |
| 44 | 1 | 33 | 0.157 | 0.030 | 0.670 | 0.225 |
| 45 | 26 | 35 | 0.166 | 0.066 | 0.275 | 0.486 |
| 46 | 8 | 35 | 0.136 | 0.110 | 0.275 | 0.353 |
| 47 | 0 | 36 | 0.196 | 0.061 | 0.196 | 0.762 |
| 48 | 2 | 36 | 0.188 | 0.080 | 0.196 | 0.683 |
| 49 | 22 | 36 | 0.148 | 0.041 | 0.196 | 0.627 |
| 50 | 24 | 4 | 0.171 | 0.082 | 0.184 | 0.643 |
| 51 | 7 | 4 | 0.180 | 0.140 | 0.184 | 0.553 |
| 52 | 22 | 5 | 0.144 | 0.041 | 0.533 | 0.251 |
| 53 | 24 | 5 | 0.187 | 0.082 | 0.533 | 0.304 |
| 54 | 26 | 5 | 0.157 | 0.066 | 0.533 | 0.261 |
| 55 | 7 | 5 | 0.230 | 0.140 | 0.533 | 0.341 |
| 56 | 0 | 6 | 0.183 | 0.061 | 0.183 | 0.750 |
| 57 | 10 | 6 | 0.164 | 0.063 | 0.183 | 0.666 |
| 58 | 24 | 7 | 0.170 | 0.082 | 0.384 | 0.364 |
| 59 | 6 | 7 | 0.130 | 0.069 | 0.384 | 0.287 |
| 60 | 10 | 8 | 0.158 | 0.063 | 0.136 | 0.791 |

**Table S4.**
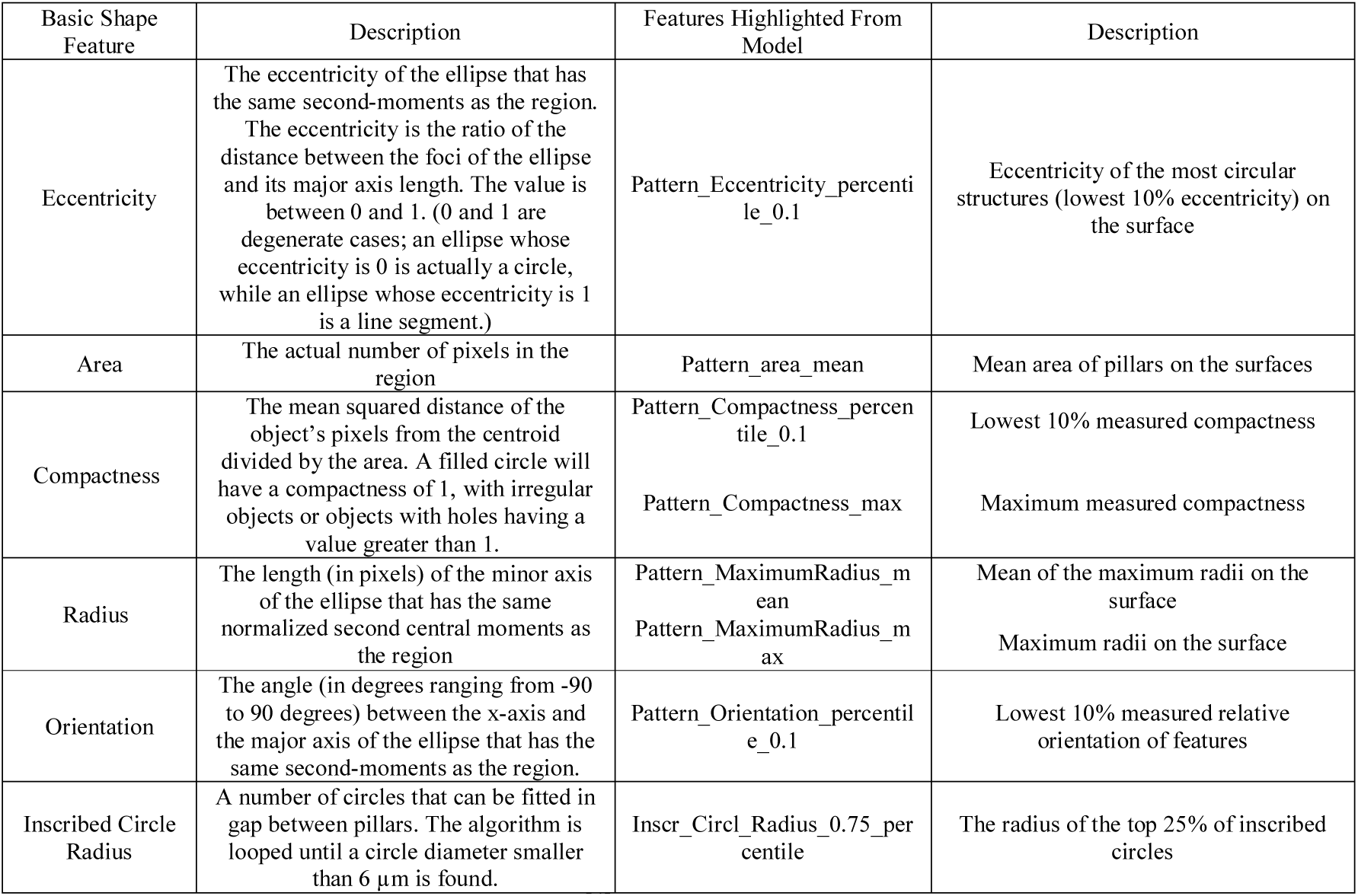
Topographical Descriptors:^[4]^.

## Notes

### Summary of Updates

The following disclosure has been added to the website and the manuscript file. Jan deBoer is a founder and shareholder of Materiomics b.v.

